# Epstein-Barr virus latency programs dynamically sensitize B-cells to ferroptosis

**DOI:** 10.1101/2022.02.21.481361

**Authors:** Eric M. Burton, Jewel Voyer, Benjamin E. Gewurz

**Affiliations:** Division of Infectious Diseases, Department of Medicine, Brigham and Women’s Hospital, 181 Longwood Avenue, Boston MA 02115, USA; Harvard Program in Virology, Boston, MA 02115; Broad Institute of Harvard and MIT, Cambridge, MA 02142, United States of America; Department of Microbiology, Harvard Medical School, Boston, MA 02115, USA

**Keywords:** lipid metabolism, oxidative stress, lipid hydroperoxide lipid metabolism, reactive oxygen species, herpesvirus, B-lymphocyte, tumor virus, ferroptosis inducing agent, erastin, ML-210, glutathione metabolism, selenocysteine, tumor suppressor, CD40, B-cell receptor, toll-like receptor, Burkitt

## Abstract

Epstein-Barr virus (EBV) causes 200,000 cancers annually. Upon B-cell infection, EBV induces lipid metabolism to support B-cell proliferation. Yet, little is known about how latent EBV infection, or human B-cell stimulation more generally, alter sensitivity to ferroptosis, a non-apoptotic form of programmed cell death driven by iron-dependent lipid peroxidation and membrane damage. To gain insights, we analyzed lipid reactive oxygen species (ROS) levels and ferroptosis vulnerability in primary human CD19+ B-cells infected by EBV or stimulated by key B-cell receptors. Prior to the first mitosis, EBV-infected cells were exquisitely sensitive to blockade of glutathione biosynthesis, a phenomenon not observed with B-cell receptor stimulation. Subsequently, EBV-mediated Burkitt-like hyper-proliferation generated elevated levels of lipid ROS, which necessitated SLC7A11-mediated cystine import and glutathione peroxidase 4 (GPX4) activity to prevent ferroptosis. By comparison, B-cells were sensitized to ferroptosis induction by combinatorial CD40-ligand and interleukin-4 stimulation or anti-B-cell receptor and Toll-like receptor 9 stimulation upon GPX4 inhibition, but not with SLC7A11 blockade. EBV transforming B-cells became progressively resistant to ferroptosis induction upon switching to the latency III program and lymphoblastoid physiology. Similarly, latency I Burkitt cells were particularly vulnerable to blockade of SLC7A11 or GPX4 or cystine withdrawal, while latency III Burkitt and lymphoblastoid cells were comparatively resistant. The selenocysteine biosynthesis kinase PSTK was newly implicated as a cellular target for ferroptosis induction including in Burkitt cells, likely due to roles in GPX4 biosynthesis. These results highlight ferroptosis as an intriguing therapeutic target for the prevention or treatment of particular EBV-driven B-cell malignancies.

**Significance:** EBV contributes to B-cell Burkitt and post-transplant lymphoproliferative disease (PTLD). EBV transforming programs activate lipid metabolism to convert B-cells into immortalized lymphoblastoid cell lines (LCL), a PTLD model. We found that stages of EBV transformation generate lipid reactive oxygen species (ROS) byproducts to varying degrees, and that a Burkitt-like phase of B-cell outgrowth is dependent on lipid ROS detoxification by glutathione peroxidase 4 and its cofactor glutathione. Perturbation of this redox defense in early stages of B-cell transformation or in Burkitt cells triggered ferroptosis, a programmed cell death pathway. LCLs were less dependent on this defense, a distinction tied to EBV latency programs. This highlights ferroptosis induction as a novel therapeutic approach for prevention or treatment of EBV+ lymphomas.

## Introduction

Epstein-Barr virus (EBV) is an oncogenic *γ*-herpesvirus that persistently infects over 95% of adults worldwide. Although typically benign, EBV contributes to >200,000 cancers/year, including endemic Burkitt, Hodgkin and non-Hodgkin lymphomas (1–3). The latter include central nervous system (CNS) lymphomas in people with advance human immunodeficiency virus infection and post-transplant lymphoproliferative diseases (PTLD). EBV is also associated with nasopharyngeal and gastric carcinoma (4–6).

EBV infects tonsillar B-cells and uses a series of latency programs to then navigate the B-cell compartment, in which different combinations of Epstein-Barr nuclear antigens (EBNA) and latent membrane protein (LMP) are expressed (7, 8). *In vitro*, EBV converts resting B-cells into immortalized lymphoblasts through a series of viral latency programs (5, 7). Over the first several days post-infection, the EBV pre-latency program is expressed, where EBNA2 drives particularly high levels of the MYC proto-oncogene (9–12). Over this stage of infection, B-cells quadruple in volume as they remodel, but do not yet divide (13, 14). Over days 4-7 post-infection, cells express Latency IIb, comprised of EBNA1, 2A and LP and non-coding RNAs (14). Latency IIb-driven cells have elevated MYC levels and undergo Burkitt-like hyper-proliferation (13, 15). Aberrant MYC expression is also a hallmark of Burkitt lymphoma, driven by chromosomal translocations between the *MYC* and immunoglobulin loci (16, 17).

EBV-infected B-cells transition to lymphoblastoid physiology at approximately one week post-infection and begin to express latency III, comprised of six EBNA and two Latent Membrane Proteins (LMP) (18). LMP1 mimics signaling by CD40, a key B-cell co-receptor activated by T-cell CD40-ligand (CD40L) (19–22). LMP2A mimics aspects of B-cell receptor signaling (23, 24). Decreasing MYC levels and likely also feedback regulation enable steadily increasing LMP1 levels over the first weeks of the lymphoblastoid phase (25). Infected cells can then be grown as immortalized lymphoblastoid cell lines (LCL), which model PTLD and CNS lymphomas, which express the EBV latency III program.

EBNA2 and MYC strongly induce host lipid metabolism (12, 15). LMP1 also induces fatty acid synthesis enzymes at later timepoints (26). EBV-induced enzymes contribute to the biosynthesis of membrane phospholipid polyunsaturated fatty acids, which are implicated in ferroptosis (27). We therefore hypothesized that the pronounced increase in lipid metabolism, together with EBV-driven induction of oxidative phosphorylation reactive oxidative species (ROS) (12, 28, 29), sensitizes newly infected B cells to ferroptosis. The recently identified non-apoptotic ferroptosis programmed cell death pathway is driven by iron-catalyzed lipid ROS, which causes irreversable membrane peroxidation(30–32). Ferroptosis sensitivity is tightly linked to amino acid metabolism, particularly of cysteine, whose availability is limiting for the biosynthesis of the major cellular antioxidant glutathione (33). Most extracellular cysteine exists in the oxidized cystine state. Therefore, cells often import cystine via the SLC7A11/xCT transporter and then reduce it to cysteine, or synthesize cysteine via the methionine metabolism transsulfuration pathway. The enzyme glutathione peroxidase 4 (GPX4) protects cells against oxidative stress by utilizing reduced glutathione (GSH) as the main reductant to resolve lipid ROS into alcohols (32).

Ferroptosis has been studied across a range of human cancer models, which show variable sensitivity to its induction (27, 34). However, effects of EBV or other tumor viruses on lipid ROS production and ferroptosis sensitization have not been studied. Likewise, effects of human primary B-cell stimulation by physiological agonists such as CD40L, cytokine stimulation, B-cell immunoglobulin receptor crosslinking or Toll-like receptor (TLR) activation are uncharacterized. Here, we investigated lipid ROS and ferroptosis sensitivity across a timecourse of primary B-cell immortalization by EBV, by B-cell receptor activation, and in EBV-transformed Burkitt and lymphoblastoid B-cells.

## Results

### EBV or receptor stimulation promote primary B cell lipid peroxidation

We hypothesized that EBV-induced lipid metabolism sensitizes newly infected B-cells to ferroptosis inducing agents. To investigate this possibility, we purified primary human CD19+ peripheral blood B cells by negative selection. B-cells were then infected by the EBV B95.8 strain at a multiplicity of infection of 0.1. We measured lipid ROS levels in uninfected cells and at 8 timepoints post-EBV infection that span B-cell transformation into LCLs (Fig. 1A) using fluorescence-activated cell sorting (FACS) analysis of the the lipid peroxidation sensor dye BODIPY-C11. EBV triggered lipid ROS production within the first two days of infection, which substantially increased upon entry into Burkitt-like hyper-proliferation between days 4-7 post-infection. Lipid ROS levels remained elevated, but progressively declined as cells converted to the lymphoblastoid phase (Fig. 1B).

**Fig. 1.**
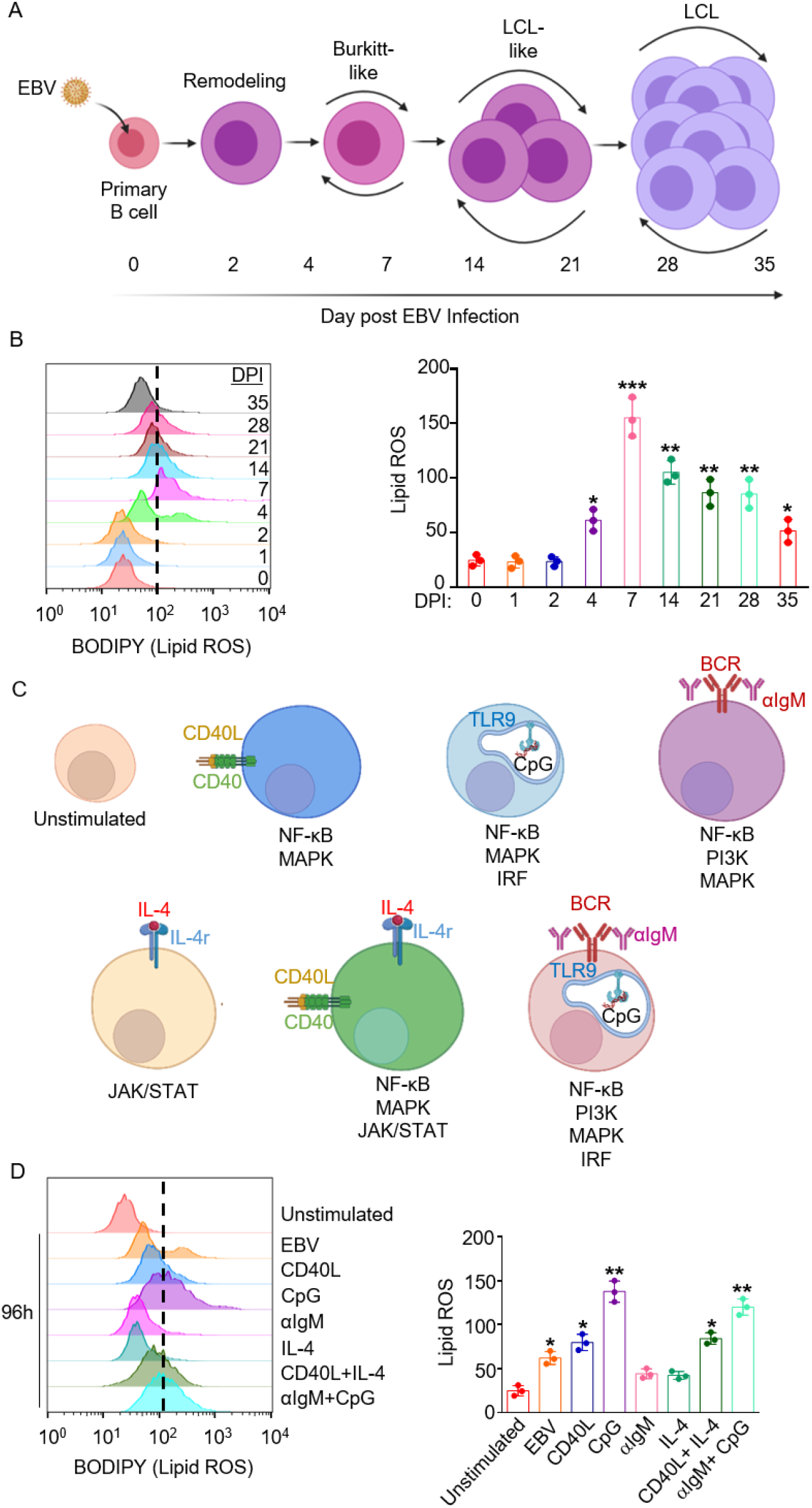
B cell EBV infection and immune receptor stimulation trigger varying levels of lipid ROS. (A) Model of the stages of EBV-mediated B-cell transformation into LCLs. (B) Representative FACS histograms of primary B-cell BODIPY (lipid ROS) levels at the indicated day post EBV-infection (DPI). Shown at right are the mean fluorescence intensity (MFI) + SD of lipid ROS levels from n=3 independent replicates. (C) Model of agonists used to stimulate primary B-cells. Key pathways activated by each mode of stimulation are shown beneath each cell model. (D) Representative FACS histograms of primary B-cell BODIPY (lipid ROS) levels at day 4 of the indicated stimulation: Mega-CD40L (50 ng/mL), αIgM (1 μg/mL), CpG (1 μM) or IL-4 (20 ng/mL). Day zero unstimulated cells were included for reference. Shown at right are the BODIPY MFI + SD values from n=3 independent replicates. P-values were determined by one-sided Fisher’s exact test. * p<0.05, **p<0.005, ***p<0.0005.

To place the above EBV effects in a broader B-cell context, we next stimulated primary B-cells by key activation pathways: CD40, interleukin 4 (IL-4), B-cell immunoglobulin crosslinking (αIgM) or toll-receptor 9 (TLR9). Notably, T-cells can activate B-cells through CD40L+IL-4 co-stimulation, whereas gain-of-function CD79 and MyD88 mutations downstream of surface immunoglobulin and TLR9 often co-occur in diffuse large B-cell lymphoma (35) (Fig 1C). After 96 hours of stimulation, lipid ROS levels were provoked by CpG, and to a somewhat lesser extent by CD40L, but not by IL-4 or αIgM treatment alone. Combinatorial CD40L+IL-4 or αIgM + CpG treatment did not induce levels beyond those seen with CD40L or CpG alone (Fig. 1D). To gain insights into how these stimuli might differentially affect lipid ROS levels in responding non-dividing vs. proliferating cells, primary B-cells were labeled by CellTrace Violet and stimulated by individual or combinatorial receptor agonists for four days. Proliferation was monitored by FACS dye dilution assay. As expected, stimulation by CD40L, IL-4, αIgM or CpG alone did not trigger robust proliferation responses, but did increase lipid ROS in the small population of proliferating cells. By contrast, cells proliferating in response to αIgM/CpG and to CD40/IL-4 had only modestly elevated lipid ROS levels as compared to the non-dividing population. These results suggest that combinatorial stimuli may buffer increases in lipid ROS levels in the proliferating cell population to facilitate outgrowth (Fig. S1A-B), and that EBV may not trigger higher lipid ROS levels than physiological B-cell activation.

### EBV dynamically sensitizes B-cells to ferroptosis

Erastin is a highly-selective SLC7A11 inhibitor (36) that induces lipid ROS and ferroptosis by restricting the supply of cysteine, the rate limiting component of glutathione biosynthesis (Fig. 2A). While erastin did not alter resting human CD19+ B cell lipid ROS or viability (Fig. S1C-E), EBV sensitized B-cells to erastin-induced death at 2 days post-infection, as judged by uptake of the vital dye 7-aminoactinomycin D (7-AAD) (Fig. 2B). Erastin sensitivity was greatest over the first week post-infection, with as many as >60% of cells positive for 7-AAD at day 7. Erastin-mediated cell death could be blocked by co-administration of the lipophilic antioxidant ferrostatin-1 (Fer-1), which scavenges lipid ROS (37). These results suggest that erastin induced ferroptosis in newly EBV-infected B-cells (Fig. 2B).

**Fig. 2.**
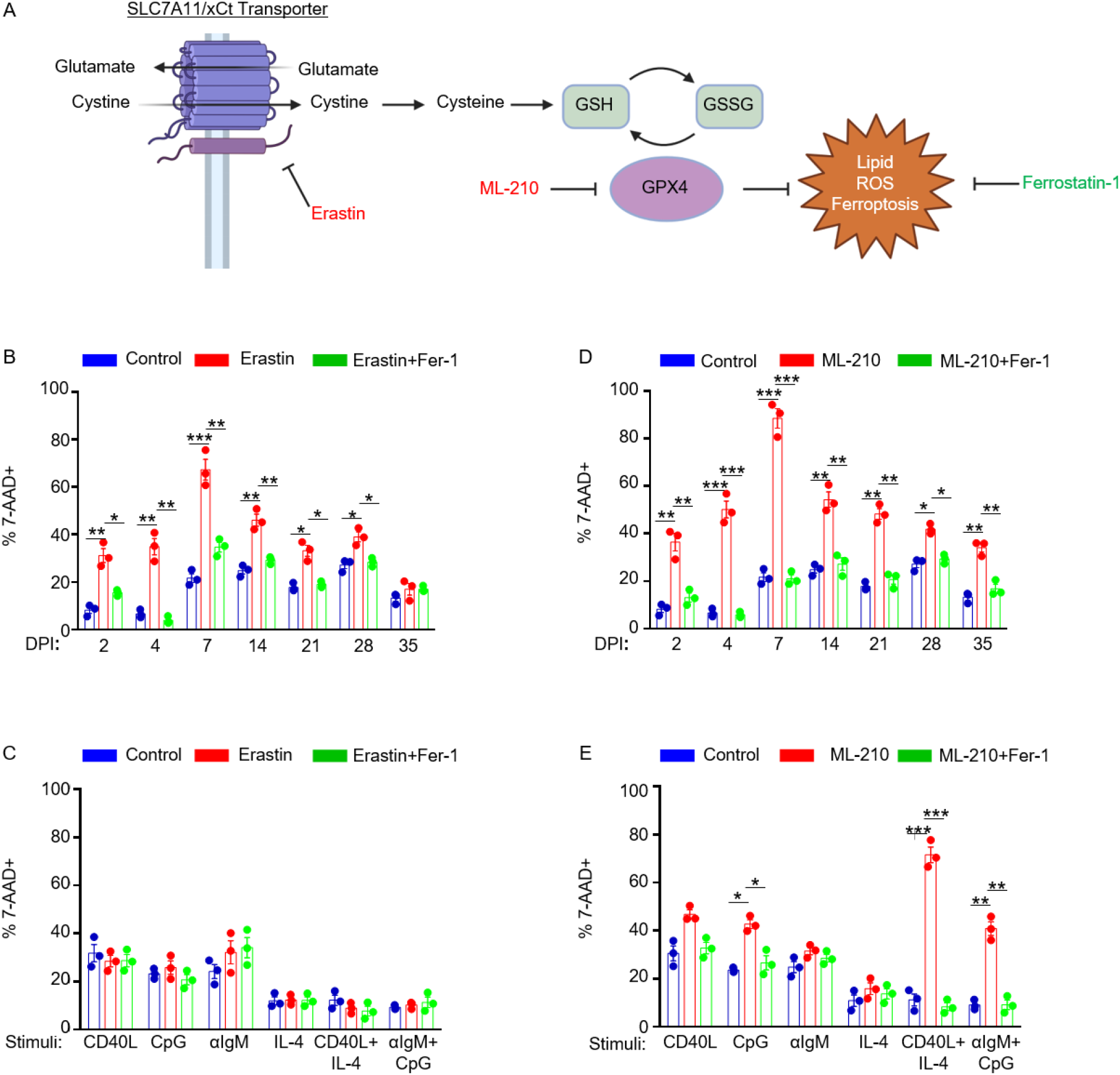
Effects of EBV infection versus B-cell receptor stimulation on sensitization to ferroptosis. (A) Model of erastin and ML-210 ferroptosis inducing agent mechanisms of action on ferroptosis induction. GSH, reduced glutathione. GSSG, oxidized glutathione. (B) FACS %7AAD+ mean + SD values from n=3 independent replicates of primary B-cells from independent donors infected by EBV for the indicated DPI and treated with 10 μM erastin and/or 5 μM Fer-1 for 24 hours prior to analysis, as indicated. (C) FACS %7AAD+ mean + SD values from n=3 independent replicates of primary B-cells from independent donors stimulated as indicated by Mega-CD40L (50 ng/mL), αIgM (1 μg/mL), CpG (1 μM) or IL-4 (20 ng/mL) and treated with 10 μM erastin and/or 5 μM Fer-1 for 24 hours prior to analysis, as indicated. (D) FACS %7AAD+ mean + SD values from n=3 independent replicates using primary B-cells from independent donors infected by EBV for the indicated DPI and treated with 1 μM ML-210 and/or 5 μM Fer-1 for 24 hours prior to analysis, as indicated P-values were determined by one-sided Fisher’s exact test. *p<0.05, ** p<0.005, *** p<0.0005.

Erastin sensitivity progressively diminished as EBV-infected cells converted to lymphoblastoid physiology (Fig. 2B and S1F) and express increasing levels of LMP1 (25). By contrast, stimulation by the panel of B-cell agonists, including combinatorial CD40+IL-4 and αIgM+CpG stimuli, failed to sensitize B-cells to erastin, despite increasing lipid ROS levels (Fig. 1D, 2C and S1G). These results suggest that B-cell activation by EBV versus immune receptor stimuli may trigger different routes of cysteine acquisition.

Whether primary human B-cells are sensitive to GPX4 loss or ferroptosis is not known. Murine follicular B-cells are not dependent on GPX4 for growth or survival, whereas marginal zone and B-1 B-cells display elevated levels of lipid metabolism and require GPX4 to prevent lipid peroxidation and ferroptosis *in vivo* (38). We therefore asked if EBV infection, or B-cell immunoreceptor stimulation more generally, confer GPX4 dependency on human B-cells. Peripheral blood CD19+ B-cells were EBV infected or stimulated by a range of B-cell stimuli and then challenged with ML-210, a potent and selective GPX4 antagonist (Fig. 2A)(39). Resting B cells were not affected by ML-210 treatment (Fig. S1B-D). However, EBV-infected B cells were vulnerable to ML-210 across the transformation time-course, reaching levels of nearly 100% 7-AAD positivity at day 7 post-infection, the timepoint with the greatest lipid ROS level (Fig. 1B, 2D, and S2A). Importantly, fer-1 treatment rescued ML-210-induced cell death, supporting on-target ML-210 effects on the ferroptosis pathway. Despite similarly robust lipid ROS induction by CpG stimulation (Fig. 1D), ML-210 more robustly induced ferroptosis in cells stimulated by combinatorial CD40L+IL-4 or by αIgM+CpG, as judged by the fold-change of 7-AAD+ cells from control or Fer-1 treated cell levels (Fig. 2E and S2B). These data suggest that the route of B-cell stimulation is a major determinant of ferroptosis vulnerability, and that human peripheral blood B-cells require GPX4 upon EBV infection or with combinatorial B-cell receptor activation.

### EBV+ Latency I Burkitt cells are more vulnerable to ferroptosis than Latency III LCLs

We next asked whether similar relationships could be observed in EBV-transformed B-cells with distinct latency states. Burkitt lymphoma (BL) is driven by aberrant MYC expression, reminiscent of newly infected cells over the first several days of infection (16). Erastin significantly increased lipid ROS and 7-AAD uptake in EBV+ Daudi BL cells, but not in the LCL GM12879 (Fig. 3A-B, S3A-B). Fer-1 blocked erastin-induced lipid ROS and cell death, suggesting on-target effects on ferroptosis induction in Daudi cells. In further support, CRISPR/Cas9 *SLC7A11* knockout (KO) significantly reduced viable cell numbers in Daudi, but not GM12878 LCLs, which could be rescued by Fer-1 (Fig. 3C-D, Table S1). Similar results were observed in another BL/LCL pair (Fig. S3A-F). These results suggest that Burkitt cells are significantly more dependent than LCLs on cystine uptake via SLC7A11 to support redox defense.

**Fig. 3.**
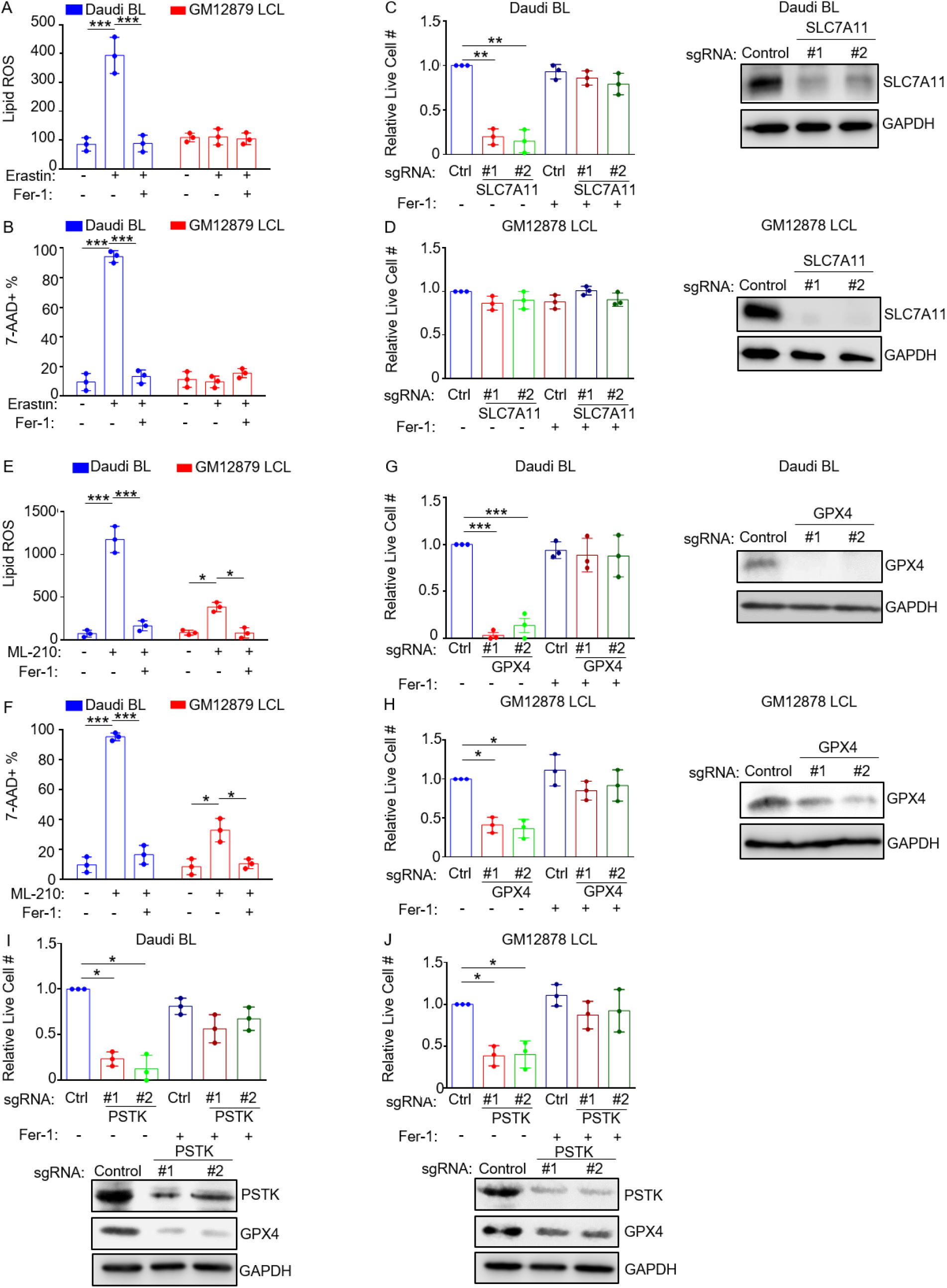
EBV infected Burkitt lymphoma cells are more susceptible to FINs than Lymphoblastoid cell lines. (A) BODIPY MFI + SD values from n=3 independent replicates of Daudi BL vs GM12879 LCLs treated with erastin (10μM) for 18 hours and measured by FACS. (B) %7AAD+ mean + SD values from n=3 replicates of Daudi and GM12879 cells treated with erastin (10 μM) for 24 hours and measured by FACS. (C-D) Relative mean + SD live cell numbers from CellTiterGlo analysis from n=3 replicates of Cas9+ Daudi (C) versus GM12878 LCLs (D) after 7-days of control (ctrl) or independent SLC7A11 sgRNA expression, in the absence or presence of 5 μM Fer-1, as indicated. Shown at right are representative immunoblots of whole cell lysates (WCL) for SLC7A11 or GAPDH load control. (E) BODIPY MFI + SD values from n=3 independent replicates of Daudi BL vs GM12879 LCLs treated with ML-210 (1μM) for 18 hours (F) 7-AAD+ mean + SD values from n=3 replicates of Daudi and GM12879 cells treated with ML-210 (1μM) for 24 hours and measured by FACS. (G and H) Relative mean + SD live cell numbers from CellTiterGlo analysis from n=3 replicates of Cas9+ Daudi (G) versus GM12878 LCLs (H) after 7-days of ctrl or GPX4 sgRNA expression, in the absence or presence of 5 μM Fer-1, as indicated. Shown at right are representative immunoblots of whole cell lysates (WCL) for GPX4 or GAPDH load control. (I and J) Relative mean + SD live cell numbers from CellTiterGlo analysis from n=3 replicates of Cas9+ Daudi (I) versus GM12878 LCLs (J) after 7-days of ctrl or PSTK sgRNA expression, in the absence or presence of 5 μM Fer-1, as indicated. Shown at right are representative immunoblots of whole cell lysates (WCL) for PSTK or GAPDH load control. P-values were determined by one-sided Fisher’s exact test. * p<0.05, **p<0.005, ***p<0.0005.

We next tested whether BLs and LCLs were dependent on GPX4 for ferroptosis resistance. Interestingly, GPX4 inactivation by ML-210 strongly induced lipid ROS and cell death in Daudi BL, but to a lesser extent in GM12879 LCLs. These phenotypes were rescued by Fer-1, suggestive of on-target effects on lipid ROS and ferroptosis (Fig. 3E-F, S4A-B). Similar results were obtained in a second BL/LCL pair (S4C-D). CRISPR *GPX4* editing by independent sgRNAs also significantly reduced BL cell survival to a somewhat greater extent than LCLs, in a manner rescuable by Fer-1 (Fig. 3G-H, Table S1, S4E-F).

GPX4 utilizes selenocysteine as a component of its catalytic triad (40). Interrogation of Cancer Dependency Map (DepMap) CRISPR data from >700 cell lines (41) demonstrated a particularly strong co-dependency between GPX4 and the kinase phosphoseryl tRNA kinase (PSTK) (Fig. S4G). PSTK phosphorylates seryl-tRNA(Sec) in order to yield O-phosphoseryl-tRNA(Sec), an activated intermediate used in selenocysteine biosynthesis (42). Since this kinase has not previously been studied in the context of ferroptosis or in EBV-infected cells, we used CRISPR to test effects of PSTK KO. We found that PSTK and GPX4 KO exhibited similar phenotypes, with more robust effects on BL than LCL survival (Fig 3I-J, Table S1, S4H-I). Intriguingly, CRISPR PSTK depletion also reduced GPX4 steady state levels, suggesting a potential PSTK role in GPX4 biosynthesis or stability (Fig 3I-J, S4H-I). Taken together, these data suggest that Burkitt cells, and to a lesser extent LCLs, are dependent on GPX4 for lipid ROS detoxification and ferroptosis defense, and that PSTK is a potentially druggable ferroptosis pathway kinase target.

### The EBV latency III induces resistance to ferroptosis inducing agents

To gain insights into how EBV latency programs alter lipid ROS levels and ferroptosis vulnerability, we utilized isogenic Burkitt tumor derived MUTU I vs III cell lines, which differ only by EBV latency I vs III programs, (43). Lipid ROS levels were somewhat higher at baseline in MUTU III than in MUTU I (Fig. 4A, S5A). Erastin treatment increased lipid ROS levels in both, although the fold change from baseline was higher in MUTU I (Fig 4A, S5A). Fer-1 restored baseline lipid ROS levels in erastin-treated MUTU I and III. Consistent with the higher fold-change in lipid ROS levels seen in MUTU I, erastin also caused higher levels of cell death in MUTU I treated for 24 hours, which could be rescued by Fer-1, indicative of ferroptosis (Fig 4B, S5B). ML-210 induced more lipid ROS in MUTU I than in MUTU III and induced higher levels of cell death, again rescuable by Fer-1 (Fig. 4CD, S5C-D). Latency III cell lines had substantially higher erastin and ML-210 half maximal inhibitory concentration (IC_50_) than latency I B-cell lines (Fig. 4E-H). For instance, the LCL Kem III had substantially higher erastin and ML-210 IC_50_ values than latency I Kem I BL cells cultured from the same tumor sample (44). This phenotype was evident in Burkitt cells harboring either of the two major EBV strain types, suggesting evolutionary conservation.

**Fig. 4:**
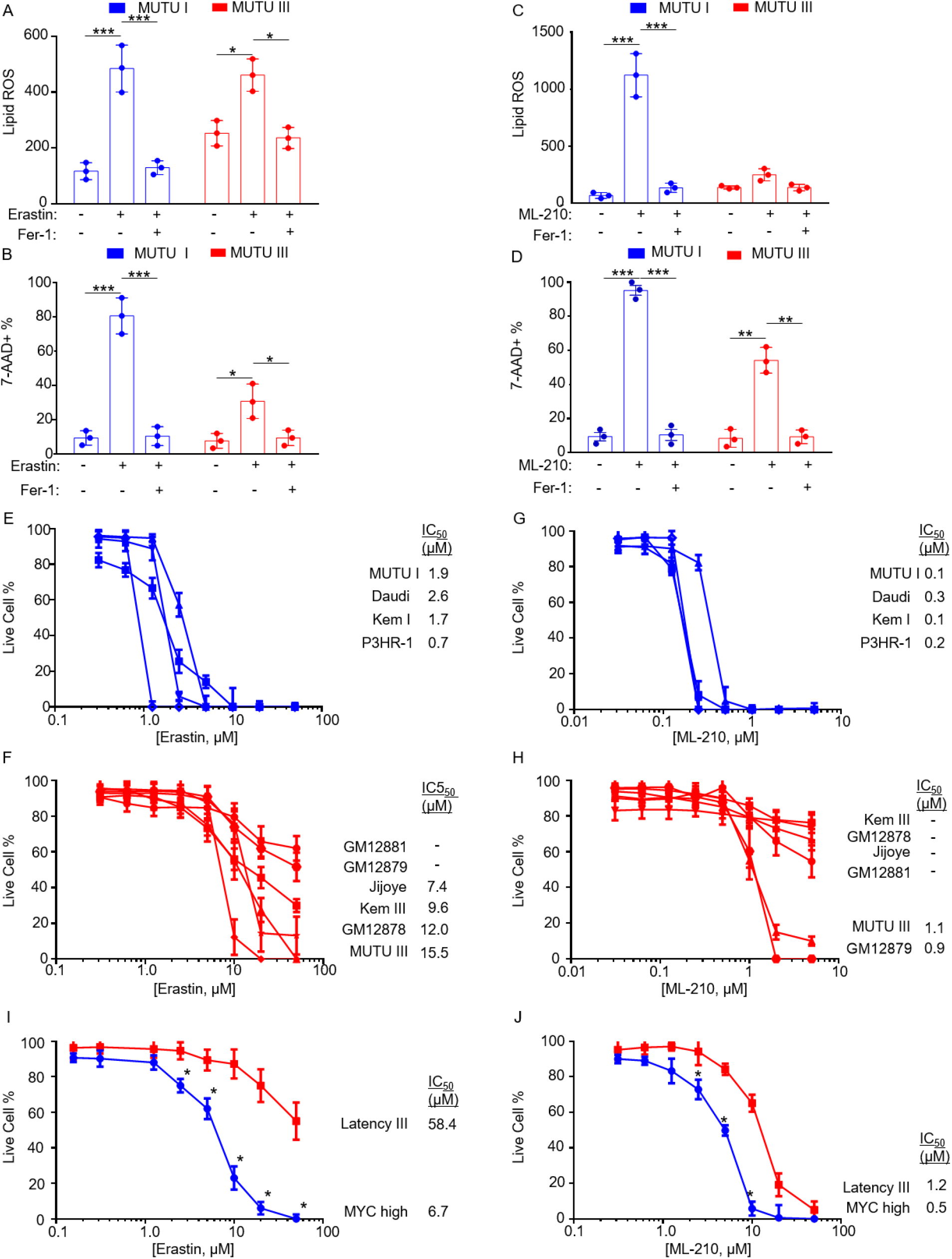
The EBV latency III program confers resistance to ferroptosis inducing agents. (A) BODIPY MFI + SD values from n=3 independent replicates of MUTU I vs III BL cells treated with erastin (10 μM) and/or Fer-1 (5 μM) for 18 hours and measured by FACS. (B) %7AAD+ mean + SD values from n=3 replicates of MUTU I vs III BL cells treated with erastin (10 μM) and/or Fer-1 (5 μM) for 24 hours and measured by FACS. (C) BODIPY MFI + SD values measured by FACS of n=3 independent replicates of MUTU I vs III BL cells treated with ML-210 (1 μM) and/or Fer-1 (5 μM) for 18 hours as indicated. (D) %7AAD+ mean + SD values measured by FACS of n=3 replicates of MUTU I vs III BL cells treated with ML-210 (1 μM) and/or Fer-1 (5 μM) for 24 hours and measured by FACS. (E) Dose-response curves of cells with EBV Latency I to erastin for 24 hours. Mean + SD of CellTiterGlo live cell numbers from n=3 replicates. IC_50_ values are shown at right. (F) Dose-response curves of cells with EBV Latency III to erastin for 24 hours. Mean + SD of CellTiterGlo live cell numbers from n=3 replicates. IC_50_ values are shown at right. (G) Dose-response curves of cells with EBV Latency I to ML-210 for 24 hours. Mean + SD of CellTiterGlo live cell numbers from n=3 replicates. IC_50_ values are shown at right. (H) Dose-response curves of cells with EBV Latency III to ML-210 for 24 hours. Mean + SD of CellTiterGlo live cell numbers from n=3 replicates. IC_50_ values are shown at right. (I) Dose-response curves of P-493 cells grown in the MYC-high Burkitt-like state (blue) or EBNA2-driven latency III state (red) to erastin for 24 hours. Mean + SD of CellTiterGlo live cell numbers from n=3 replicates. IC_50_ values are shown at right. (J) Dose-response curves of P-493 cells grown in the MYC-high Burkitt-like state (blue) or EBNA2-driven latency III state (red) to ML-210 for 24 hours. Mean + SD of CellTiterGlo live cell numbers from n=3 replicates. IC_50_ values are shown at right. P-values were determined by one-sided Fisher’s exact test. * p<0.05, **p<0.005, ***p<0.0005.

We next asked whether switching off the latency III program in an LCL, and replacing it with conditional MYC over-expression in the context of latency I, altered sensitivity to ferroptosis. We utilized the P493-6 LCL, which have conditional Tet-off MYC and 4-hydroxytamoxifen (4HT) activated EBNA2 alleles (45). P493-6 grown in 4HT and doxycycline grow as LCLs, whereas cells grown without either lose latency III and express high levels of MYC to simulate EBV+ BL cell physiology (45)(Fig. S5E). P493-6 grown in the LCL state exhibited significantly higher erastin IC_50_ values (58.4 vs 6.7 μM) and likewise for ML-210 (1.2 vs 0.5 μM) (Fig. 4I-J). These results suggest that MYC-driven lipid metabolism in newly EBV-infected and Burkitt cells induces dependency on SLC7A11 cystine import, glutathione biosynthesis and GPX4-based ferroptosis defense.

### Glutathione biosynthesis is critical for Burkitt-like and Burkitt cell survival

The enzyme glutamate-cysteine ligase (GCLC) catalyzes the rate limiting step in glutathione biosynthesis, the production of *γ*-glutamylcysteine from cysteine and glutamate (Fig. 5A). Given the observed differences in EBV-infected cell sensitivity to blockade of SLC7A11 or GPX4, as well as our prior observation that resting B-cells have low levels of glutathione (12), we next asked whether newly EBV-infected cells are dependent on glutathione synthesis for survival. We treated CD19+ primary B-cells at 7 timepoints post-infection with buthionine sulfoximine (BSO), a highly selective GCLC antagonist. We then monitored effects on 7-AAD uptake and lipid ROS levels. Unexpectedly, cells were exquisitely sensitive to BSO treatment at Day 2 post-EBV infection, where we observed highly elevated lipid ROS levels and >80% underwent cell death (Fig. 5B-C, and Fig S6A-B). Supplementation with either Fer-1 or GSH reduced lipid ROS and significantly rescued viability, suggesting that BSO induced ferroptosis at this early timepoint post-infection (Fig. 5B-C, and Fig. S6A-B). Surprisingly, we did not observe significant BSO-driven increases in lipid ROS or cell death at later times (Fig. 5A-B). It is possible that BSO did not completely block GCLC activity at the dose used. Therefore, these results are consistent with a model in which particularly low GSH stores in resting B-cells render cells particularly sensitive to GCLC inhibition at Day 2 post-infection.

**Fig. 5.**
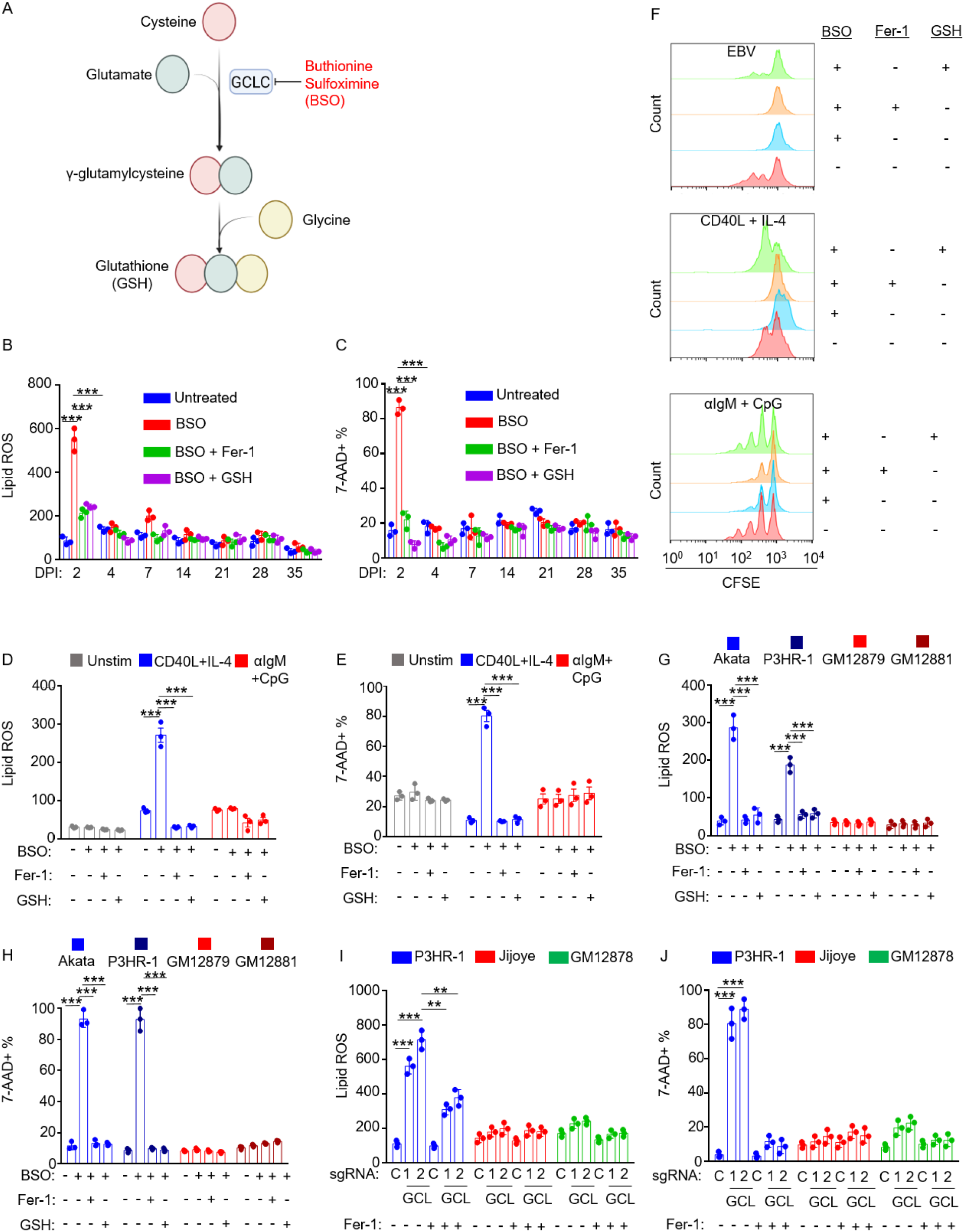
Effects of glutathione synthesis inhibition on EBV-infected, receptor stimulated and EBV-transformed B-cells. (A) Diagram of the glutathione synthesis pathway and its inhibition by BSO. (B) FACS BODIPY MFI + SD values from n=3 replicates of primary B-cells treated at the indicated day post-EBV infection with BSO (100 μM) and Fer-1 (5 μM) or 2.5mM GSH for 72 hours, as indicated.(C) %7AAD+ mean + SD values from n=3 replicates of primary B-cells treated at the indicated day post-EBV infection with BSO (100 μM) and Fer-1 (5 μM) or 2.5mM GSH for 72 hours, as indicated. (C) FACS %7-AAD+ mean + SD values from n=3 independent replicates of primary B-cells from independent donors treated at indicated day post-EBV infection with BSO (100 μM) and Fer-1 (5 μM) or 2.5mM GSH for 72 hours, as indicated (D) FACS BODIPY MFI + SD values from n=3 independent replicates of primary B-cells from independent donors stimulated for 2 days by CD40L (50 ng/mL) and IL-4 (20 ng/mL) or αIgM (1 μg/mL) and CpG (1 μM) and then treated with BSO (100 μM), 5 μM Fer-1 or 2.5 mM GSH for 48 hours prior to analysis, as indicated. (E) FACS %7-AAD+ mean + SD values from n=3 independent replicates of primary B-cells from independent donors stimulated for 2 days by CD40L (50 ng/mL) and IL-4 (20 ng/mL) or αIgM (1 μg/mL) and CpG (1 μM) and then treated with BSO (100 μM), 5 μM Fer-1 or 2.5 mM GSH for 48 hours prior to analysis, as indicated. (F) FACS analysis of CFSE-labelled primary B cells infected with EBV or treated with the indicated stimuli in the presence of BSO, Fer-1 or GSH, as indicated. After 4 days, proliferation was analyzed via FACS. Plot is representative of n=3 independent replicates. (G) FACS BODIPY MFI + SD values from n=3 independent replicates of EBV-Akata and EBV+ P3HR-1 BL or GM12879 and GM12881 LCLs treated with BSO (100 μM), Fer-1 (5 μM) or GSH (2.5 mM) for 48 hours, as indicated. (H) FACS %7-AAD+ mean + SD values from n=3 independent replicates of EBV-Akata and EBV+ P3HR-1 BL or GM12879 and GM12881 LCLs treated with BSO (100 μM), Fer-1 (5 μM) or GSH (2.5 mM) for 72 hours, as indicated. (I). FACS BODIPY MFI + SD values from n=3 independent replicates of Cas9+ latency I P3HR-1, latency III Jijoye or GM12878 LCLs expressing control (c) or GCLC (GCL) sgRNAs for 12 days and treated with Fer-1 (5 μM), as indicated. (J) FACS %7-AAD+ mean + SD values from n=3 independent replicates as in (I), analyzed 14 days after sgRNA expression. P-values were determined by one-sided Fisher’s exact test. * p<0.05, **p<0.005, ***p<0.0005.

We next cross-compared BSO effects on resting or receptor-stimulated human B-cells. BSO did not increase lipid ROS or death in resting B-cells, but strongly induced lipid ROS and 7-AAD uptake in cells stimulated for 2 days by CD40L+IL-4. Interestingly, these effects were not seen with αIgM+CpG stimulation (Fig. 5D-E, S6C-D), suggesting that the route of primary B-cell stimulation dictates dependency on GCLC enzymatic activity at early timepoints. To then measure effects on proliferation, B-cells were labeled by carboxyfluorescein succinimidyl ester (CFSE) and infected by EBV or stimulated with CD40L+IL-4 or αIgM + CpG for four days. Proliferation was monitored by FACS dye dilution assay. BSO blocked proliferation of EBV-infected and CD40+IL-4 stimulated cells, but only partially blocked αIgM+CpG-driven proliferation. BSO effects were rescued by reduced glutathione (GSH) but not Fer-1, suggesting glutathione roles beyond lipid ROS detoxification in support of B-cell proliferation (Fig. 5E).

We next examined effects of BSO treatment on a panel of Burkitt versus LCLs. BSO significantly increased lipid ROS and 7-AAD uptake in EBV+ BLs, but not LCLs (Fig. 5G-H, Fig. S6E-F). Fer-1 or GSH supplementation rescued lipid ROS and suppressed BL cell death, indicating that BSO induced ferroptosis in these BL cells. To then examine a potential relationship between EBV latency program and glutathione synthesis, we used CRISPR to deplete GCLC (Fig. S7A, Table S1). In agreement with our BSO data, *GCLC* editing markedly increased lipid ROS and cell death in P3HR-1 (Fig. 5H-I, S7B-C). By contrast, GCLC depletion in Jijoye, the parental latency III cell line of P3HR-1, or in GM12878 LCLs had little effect on lipid ROS or survival (Fig. 5H-I). Similar results were obtained with BSO treatment (Fig. S7D-F). These data suggest that latency III generates lower lipid ROS levels, utilizes distinct glutathione metabolism, and/or compensatory mechanisms to control lipid ROS.

### EBV infected and stimulated primary B cells are addicted to cystine

We next investigated whether EBV infection or B-cell stimulation renders primary or EBV-transformed B-cells auxotrophic for cystine (Fig. 6A). While cysteine is not an essential amino acid for many cell types and can be synthesized from methionine via the transsulfuration pathway, some cell states require cystine import to prevent ferroptosis (46). We therefore investigated the role of cystine restriction on B-cell lipid ROS and ferroptosis. At each of seven timepoints post-EBV infection, cystine withdrawal induced B-cell death, as judged by 7-AAD uptake (Fig. 6B). Cell viability was significantly restored by addition of Fer-1, indicating that EBV-infected B-cells require cystine to prevent ferroptosis induction. Similarly, GSH supplementation rescued cell death, perhaps as it can be metabolized extracellularly to provide cysteine. We next cross-compared effects of cystine restriction on B-cells infected by EBV versus stimulated by CD40L+IL4 or αIgM+CpG for four days. In all cases, cystine withdrawal restrained B-cell proliferation (Fig. 6C). GSH, but not Fer-1 nearly completely rescued proliferation driven by EBV, αIgM+CpG and by CD40L+IL-4 stimulation (Fig. 6C). Furthermore, whereas cystine withdrawal did not trigger lipid ROS or cell death in unstimulated B-cells, it significantly elevated lipid ROS and 7-AAD uptake in CD40L+IL-4 stimulated cells, and to a much lesser extent in αIgM+CpG treated cells (Fig. 6D-E, S8A). These results suggest that lipid metabolism programs driven by EBV infection or CD40L+IL-4 stimulation necessitate a greater dependence on cystine uptake to buffer lipid ROS.

**Fig. 6.**
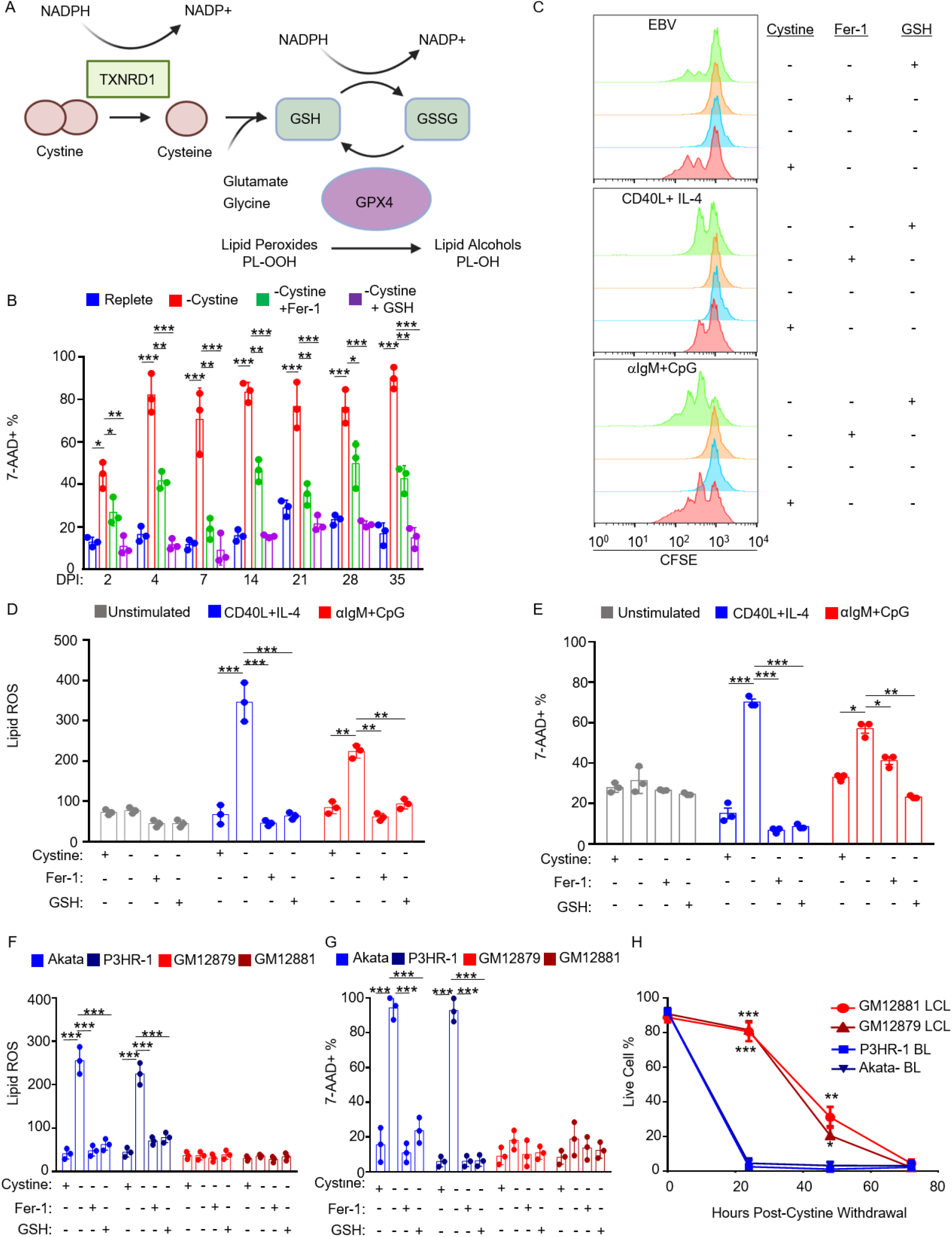
EBV infected cells are dependent on cystine for growth and survival. (A) Schematic of cystine and glutathione metabolism in control of lipid peroxidation. (B) FACS %7-AAD+ mean + SD values from n=3 independent replicates of primary B cells infected with EBV for the indicated DPI, harvested and seeded into media with or without cystine and treated with Fer-1 or GSH as indicated for 72 hours. (C) FACS analysis of CFSE-labelled primary B cells infected with EBV or treated with the indicated stimuli in the presence of Fer-1 or GSH, as indicated. After 4 days, proliferation was analyzed via FACS. Plot is representative of n=3 independent replicates. (D) FACS BODIPY mean + SD values from n=3 replicates of primary B cells unstimulated or treated with indicated stimuli for 48 hours and then seeded into media with the same stimuli, with or without cystine, Fer-1 and GSH as indicated for 48 hours. (E) FACS %7-AAD+ mean + SD values from n=3 replicates of primary B cells as in (D) and seeded into media with or without cystine, Fer-1 and GSH as indicated for 72 hours. (F-G) FACS BODIPY mean + SD (F) and %7-AAD+ mean + SD values from n=3 replicates of Akata or P3HR-1 BL and GM12879 or GM12881 LCLs grown with or without cystine, Fer-1 and GSH as indicated for 24 hours. (H) Relative % live cells determined by CellTiter Glo assay from n=3 replicates of Akata or P3HR-1 BL and GM12879 or GM12881 LCLs grown without cystine across 72 hours. P-values were determined by one-sided Fisher’s exact test. * p<0.05, **p<0.005, ***p<0.0005.

Finally, we investigated whether extracellular cystine had similarly important roles during EBV transformed B-cell lipid ROS generation and ferroptosis defense. Cystine withdrawal significantly elevated lipid ROS levels and cell death in BL, but not in either LCL over 24 hours, rescued by Fer-1 or GSH (Fig. 6F-G, S8B-C). While able to survive for longer periods of cystine withdrawal, cystine restriction ultimately triggered LCL death, similar to what we observed when culturing day 35 post infection primary B cells in cystine restricted media (Fig. 6B, 6H, and S8D). These results are consistent with a model in which greater GSH stores enable LCLs to survive cystine withdrawal for longer periods of time than Burkitt cells. Given their resistance to even prolonged erastin treatment or SLC7A11 KO, these results also suggest that latency III cells have alternative means of meeting cysteine demand, and further suggest that they may utilize non-GPX4 pathways for lipid ROS detoxification (Fig. 7).

**Fig. 7.**
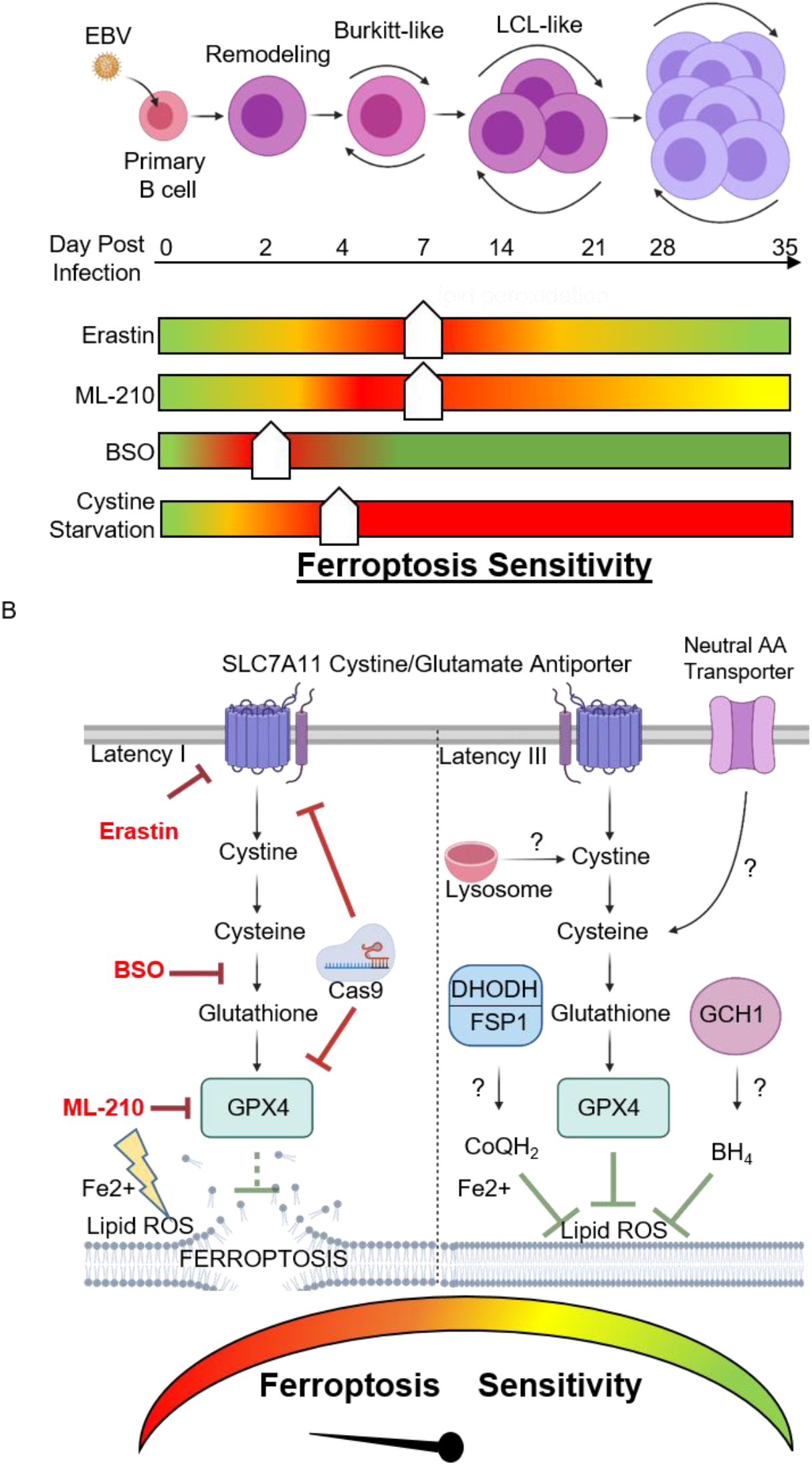
Model of EBV effects on B-cell vulnerability to ferroptosis. (A) Schematic of vulnerability to ferroptosis induction at distinct states of EBV-mediated B-cell transformation. Ferroptometer sliders demarcate the relative sensitivity of EBV-infected cells to ferroptosis induction to the indicated agent, with green and red indicating the lowest and highest sensitivities to ferroptosis induction, respectively. (B) Schematic of vulnerability to ferroptosis inducing agents in latency I (left) vs III (right) EBV-transformed B-cells. Question marks indicate potential pathways by which latency III cells may differentially acquire cysteine and buffer lipid ROS for redox defense.

## Discussion

Vulnerability to ferroptosis is associated with numerous aspects of cellular physiology, including amino acid, iron, lipid, and NADPH metabolism (32). Murine marginal zone and B1-type B cells require GPX4 to prevent ferroptosis, while murine follicular B cells do not, highlighting B-cell differentiation specific roles (38). Notably, murine B-cells have low levels of SLC7A11 expression and therefore require reducing agents such as ß-mercaptoethanol or sulfhydryl-containing compounds to survive *in vitro* (47). Human B-cells do not require addition of these reducing agents and exhibited low levels of lipid ROS at baseline. How untransformed human B-cells respond to lipid oxidative stress resulting from EBV oncogenic challenge, and how this cross-compares with B-cell receptor driven metabolic reprogramming, have remained important open questions. Here, we present evidence that latent EBV triggered lipid ROS production in newly infected primary human B-cells, which induced vulnerability to ferroptosis inducing agents at multiple times post-infection (Fig. 7). Likewise, we found that latency I Burkitt cells were exquisitely sensitive to ferroptosis induction, while latency III B-cells were more refractory (Fig. 7).

Metabolic reprogramming is required upon B-cell challenge by EBV or by receptor-driven proliferation to meet anabolic demands (12, 48, 49). Yet, metabolic stress is a major barrier to EBV-mediated B-cell transformation (28). Our results suggest that EBV must carefully balance highly induced anabolic lipid metabolism with ferroptosis defense, particularly at early timepoints post-infection and in Burkitt cells, which share hyperactive MYC states. Notably, resting human B-cells and Burkitt cells also have very low levels of glutathione and glutathione synthesis (12), in contrast to many cellular environments, where glutathione is present at millimolar quantities. To meet demand, we hypothesize that EBNA2 and MYC highly induce one-carbon metabolism, which supports glutathione biosynthesis via the production of NADPH reducing units and the glycine building block. Indeed, the majority of glutathione synthesized by day 4 post-infection could be traced through EBV-induced mitochondrial one-carbon metabolism (12). Furthermore, human B-cells also have higher ratios of oxidized to reduced glutathione than hepatocytes or erythrocytes (50). These results support a model in which EBNA2 and MYC coordinate a redox defense program, increasing one-carbon metabolism to support glutathione biosynthesis needed to suppress ferroptosis, while also providing carbon units for *de novo* purine synthesis (12) (Fig. 7).

EBV induced remarkable sensitivity to BSO at Day 2 post-infection, a phenomenon that we did not observe at later timepoints or with other forms of primary human B-cell stimulation. By contrast, many cell types and even cancer cells are resistant to BSO treatment or cystine restriction (46). Yet, EBV-infected cells were sensitive to SLC7A11 and GPX4 blockade at Day 2 and also at multiple later timepoints. We suspect that BSO engaged its GCLC target, as it highly induced lipid ROS at Day 2, which was suppressed by Fer-1 or GSH supplementation. We favor the hypothesis that Day 2 cells are exquisitely sensitive to BSO as viral transformation converts quiescent B-cells into a highly metabolically active state, in preparation for Burkitt-like B-cell hyper-proliferation. Notably, EBNA2 and EBNA-LP are expressed at particularly high levels at day 2 post-infection, which in turn hyper-induce MYC (11, 12, 51). MYC levels are higher in newly EBV-infected cells and in Burkitt cells than in B-cells stimulated by receptor agonists or in LCLs. Such aberrantly and rapidly elevated levels of EBNA2 and MYC may therefore create the acute BSO vulnerability at this timepoint, as B-cell metabolism swiftly transitions from a quiescent to a highly activated state. It is also possible that BSO did not completely block GCLC activity at the dose used, and that even a partial inhibition at Day 2 at a dose near the BSO IC_50_ was sufficient to trigger the observed massive lipid ROS and ferroptosis phenotypes, given the supply/demand imbalance at this transition point. At later timepoints, we speculate that there is sufficient glutathione reserve for the cells to buffer BSO effects over the relatively short time of BSO treatment used in our assay, and that we therefore did not observe increased lipid ROS or cell death with BSO treatment (Fig. 5B-C).

We also note that glutathione has important roles in iron-sulfur [Fe-S] cluster biogenesis. [Fe-S] clusters are highly sensitive to inactivation by ROS and play key roles in mitochondrial metabolism, including in the tricarboxylic acid cycle and oxidative phosphorylation (OXPHOS). The need to rapidly synthesize [Fe-S] clusters at Day 2 post-infection may render this timepoint particularly sensitive to BSO treatment. The resulting perturbation to [Fe-S] clusters, TCA and OXPHOS may cause reductive stress, adding to cytotoxic effects at this timepoint.

Profiling studies have revealed a wide-range of susceptibility to ferroptosis induction across human cancer cell lines, including B-cell lymphomas, yet the basis for this important observation remains to be fully elucidated (30). It is therefore notable that Burkitt cells have low glutathione stores and limited capacity for cystine uptake (52), which may render them exquisitely sensitive to ferroptosis inducing agents. Our results suggest that glutathione abundance may link the observed sensitivities to SLC7A11 and GCLC inhibition in newly-infected and Burkitt cells. Interestingly, we observed that latency III alters ferroptosis sensitivity even within the same cell context. Latency III MUTU and Jijoye Burkitt cells were more refractory to erastin or ML-210 than their latency I counterparts, and P493-6 cells were more resistant to these agents when grown in the Burkitt-like than in the LCL state. Likewise, we found that EBV-infected cells became more refractory to these agents at later timepoints of infection as primary cells undergo lymphoblastoid transformation and express latency III. We speculate that latency III-driven increases in glutathione reservoirs contribute to these phenotypes.

Latency III may further alter sensitivity to ferroptosis through additional possible mechanisms (**Fig. 7**). In particular, Latency III may utilize recently described glutathione-independent systems that detoxify lipid ROS in parallel with GPX4 (53). For instance, the oxidoreductase ferroptosis suppressor protein 1 (FSP1, also called AIFM2) reduces ubiquinone (also called coenzyme Q or CoQ) to ubiquinol (CoQH_2_) at the plasma membrane. CoQH_2_ can then trap and detoxify lipid peroxyl radicals independently of GPX4. Similarly, the *de novo* pyrimidine synthesis enzyme dihydroorotate dehydrogenase (DHODH) generates CoQH_2_, which can be used to quench lipid ROS within the mitochondrial inner membrane (54). We recently identified that EBV highly induces DHODH and de novo pyrimidine activity in newly infected B-cells (55). Furthermore, GTP cyclohydrolase 1 (GCH1) can suppress ferroptosis via the generation of the radical-trapping antioxidant tetrahydrobiopterin (BH4) (56). Use of these ferroptosis suppressing pathways could potentially explain decreased latency III cell sensitivity to SLC7A11, GCLC and GPX4 inhibition. It will therefore be of interest to test putative ferroptosis resistance roles of these recently described pathways in latency III cells.

Cysteine availability is the rate-limiting step in glutathione biosynthesis (57). Cystine is synthesized and released by the liver and is present at ~50 μM extracellular fluid. It can then be imported by SLC7A11, whereas cysteine, which is present at low concentrations extracellularly, can be imported via neutral amino-acid transporters. Our results suggest that the route of B-cell stimulation or transformation dictate specific roles of cystine/cysteine import, glutathione synthesis and GPX4. RNAseq data suggest that SLC7A11 is already upregulated by 2 days post-EBV infection, and reaches progressively higher levels at later times as cells transform to the lymphoblastoid physiology (11, 58). Thus, EBV may increase demand for cystine import through SLC7A11 particularly during Burkitt-like hyperproliferation, where lipid biosynthetic pathways may be maximally induced. We suspect that particularly at later timepoints, EBV-infected primary B-cells may also utilize other neutral amino acid transporters to acquire cysteine. For instance, the Alanine-Serine-Cysteine Transporters 1 and 2 (ASCT1 and 2) are highly induced by EBV (12). Autophagy also has a critical role in early EBV-driven B-cell hyperproliferation in the context of metabolic stress (28). Latency III cells may utilize autophagy and perhaps also pinocytosis to acquire lysosomal cysteine, which can then be pumped into the cytosol via the cystinosin lysosomal membrane transporter. Additionally, latency III may increase *de novo* cysteine synthesis through the transsulfuration pathway, reducing the demand for imported cystine.

We observed significant increases in cell death upon erastin treatment through day 28 post-infection (Fig. 2B), albeit with greatest magnitude within the first week of infection. We speculate that sensitivity is greatest at earlier timepoints where cells must import cystine to rapidly synthesize GSH and meet demand as discussed above. By contrast, we observed robust induction of cell death upon cystine withdrawal at nearly all timepoints in Fig. 6B, which was partially rescuable by Fer-1. Withdrawal of cystine from the media blocks all routes of import, and thereby may affects viability at all timepoints. It is also notable that cystine withdrawal affects the extracellular and intracellular redox states, and that this could have contributed to the observed effects on cell viability in a manner distinct from erastin effects on cystine import. Effects on redox states may also impair parallel redox defense pathways, including the antioxidant FSP1, DHODH and GCH1 pathways, thereby potentially explaining the stronger effect of cystine withdrawal than erastin at late timepoints of EBV B-cell infection and over prolonged periods in LCLs.

Current approaches for Burkitt lymphoma therapy rely on high intensity chemotherapy and cause considerable side-effects. Therefore, leveraging ferroptosis tumor suppressor roles may offer a promising approach novel approach. Encouragingly, little acute toxicity has been observed in mouse models with ferroptosis inducing agents (59). Lower cellular glutathione concentrations increases the cytotoxicity of cyclophosphamide (60), which is used in chemotherapy regimens for Burkitt lymphoma. It would be of interest to determine whether erastin or ML-210 are synergistic with cyclophosphamide in Burkitt cells. Given the sensitivity of newly-infected B-cells during Burkitt-like hyperproliferation to ferroptosis inducing agents, such strategies might also prove useful in preventing the polyclonal outgrowth of newly-EBV infected B-cells during high risk periods, including post-organ transplantation into EBV seronegative hosts, where high rates of post-transplant lymphoproliferative disease are a major clinical challenge. While GPX4 inhibitors suitable for human translational approaches remain to be developed, our results highlight the kinase PSTK as a novel potentially druggable target for the development of novel ferroptosis inducing agents.

## Materials and Methods

### B95.8 EBV preparation, primary B-cell infection and stimulation

B95-8 EBV was prepared as described (61, 62). EBV titer was determined by transformation assay. Freshly isolated, de-identified, discarded CD19+ peripheral blood B-cells obtained under our hospital institution review board approved protocol were seeded in RPMI 1640 media supplemented with 10% FBS (1 million cells/mL) for infection or stimulation studies. An EBV multiplicity of infection of 0.1 was used, or cells were stimulated by: Mega CD40L (50 ng/mL), αIgM (1 μg/mL), CpG (1 μM) and IL-4 (20 ng/mL). For 96 hour stimulation, agonists were refreshed at 48 hours. Cells were cultured in a humidified chamber at 37°C in RPMI/10% FBS.

### Primary B-cell isolation and culture

RosetteSep and EasySep negative isolation kits (Stemcell Technologies) were used sequentially to isolate CD19+ B-cells by negative selection, with the following modifications made to the manufacturer’s protocols. For RosetteSep, 40 μL of antibody cocktail was added per mL of blood and before Lymphoprep density medium was underlayed prior to centrifugation. For EasySep, 10 μL of antibody cocktail was added per mL of B cells, followed by 15 μL of magnetic bead suspension per mL of B cells. After negative selection, the cells obtained were routinely ≥95% positive for CD19, a nearly pan-B cell surface marker (though not strongly expressed on plasma cells).

### CellTiter Glo

CellTiter Glo viability assay (Promega) was performed by collecting cells at indicated timepoints and washing once with 1x PBS. 50 μL of cells in PBS were used per assay according to manufacturer instruction.

### Software/data presentation

Graphs were made using Graphpad Prism 7. Schematic models were made using Biorender.

## Acknowledgements

We would like to thank Drs. Vamsi Mootha and Thomas Sommermann for helpful discussions and suggestions. This work was supported by NIH RO1 AI137337, AI164709 and CA228700 and a Burroughs Wellcome Career Award in Medical Sciences to BEG, by T32AI007245 to E.B. and by a MARC U*Star fellowship to J.V. We thank the Harvard SHURP program for support of JV. **This work has not been peer-reviewed**.

## Author Contributions

E.B. and J.V. performed the experiments. E.B. and B.E.G. designed the study, which B.E.G supervised. E.B. and B.E.G wrote the manuscript.

