## Supplemental Figures and Legends for "Epstein-Barr virus latency programs dynamically sensitize B-cells to ferroptosis"

### Supplementary Information

#### Supplementary Figure and Legends

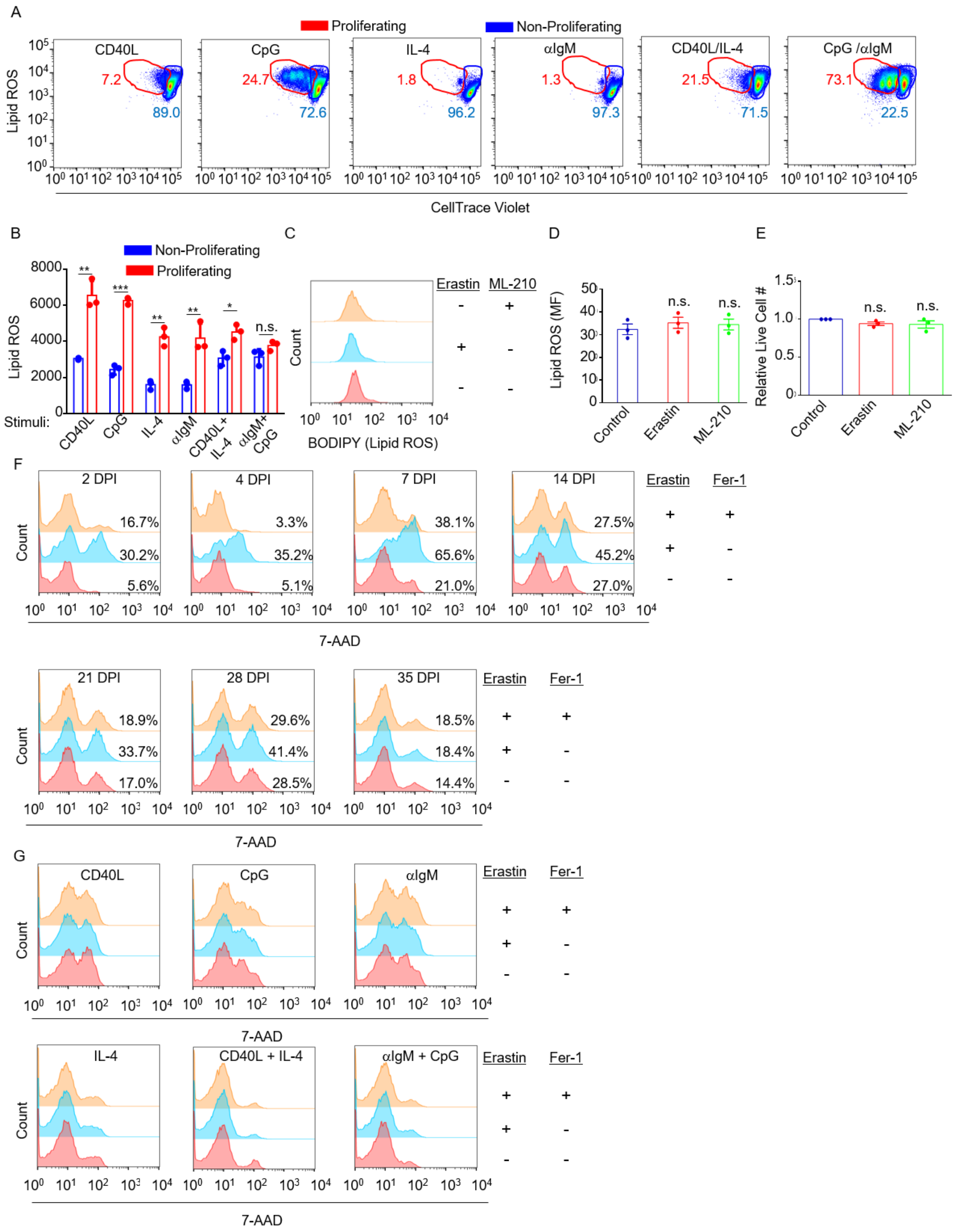

**Fig. S1: Sensitivity of EBV infected or stimulated primary B cells to ferroptosis inducing agents.**

(A) Representative FACS histograms of CellTrace Violet proliferation versus BODIPY lipid ROS staining from n=3 replicates of primary B-cells treated with Mega-CD40L (50 ng/mL), αIgM (1 μg/mL), CpG (1 μM) or IL-4 (20 ng/mL) as indicated for 4 days.

(B) Representative FACS histograms of primary B-cell BODIPY levels in resting primary human peripheral blood CD19+ B cells treated with erastin (10 μM) or ML-210 (1 μM) for 24 hours.

(C) FACS mean + SEM BODIPY lipid ROS levels from n=3 replicates as in (A).

(D) Mean + SEM relative live cell numbers from n=3 replicates as in (A), measured by CellTitreGlo.

(E) Representative FACS histograms from n=3 replicates of primary B-cell 7-AAD levels. At the indicated DPI, cells were treated with erastin (10 μM) or Fer-1 (5 μM) for an additional 24 hours, as indicated.

(F) Representative FACS histograms of 7-AAD levels from n=3 replicates of primary B-cells treated with Mega-CD40L (50 ng/mL), αIgM (1μg/mL), CpG (1 μM) or IL-4 (20ng/mL) as indicated for 3 days, with addition of erastin (10 μM) and Fer-1 (5 μM) for 24 hours, as indicated.

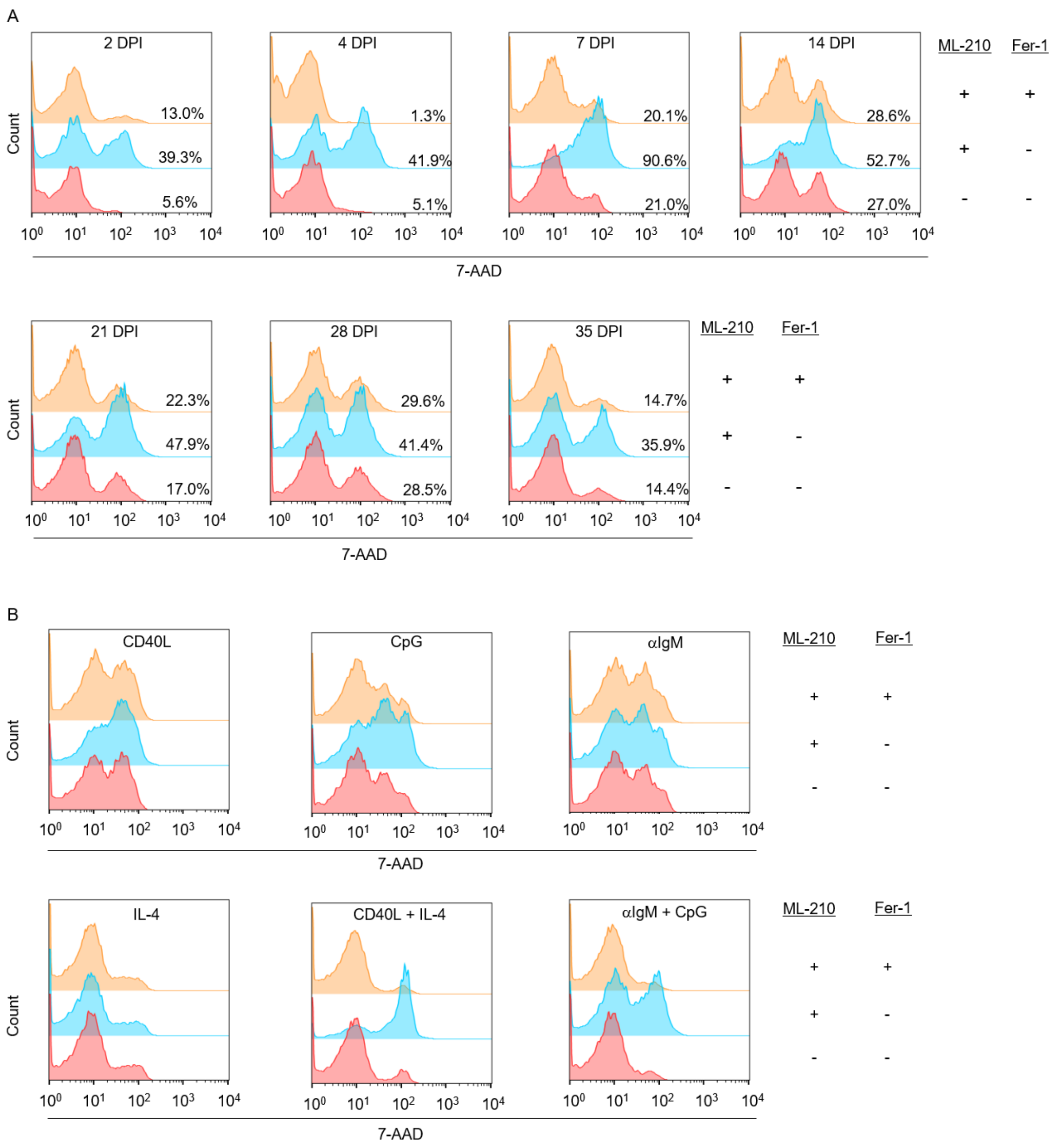


**Fig. S2: Susceptibility of EBV-infected versus stimulated B-cells to ferroptosis induction.**

(A) Representative FACS histograms from n=3 replicates of primary B-cell 7-AAD levels. At the indicated DPI, cells were treated with ML-210 (1 μM) or Fer-1 (5 μM) for an additional 24 hours, as indicated.

(B) Representative FACS histograms of 7-AAD levels from n=3 replicates of primary B-cells treated with Mega-CD40L (50 ng/mL), αIgM (1 μg/mL), CpG (1 μM) or IL-4 (20 ng/mL) as indicated for 3 days, with addition of ML-210 (1 μM) and Fer-1 (5 μM) for 24 hours, as indicated.

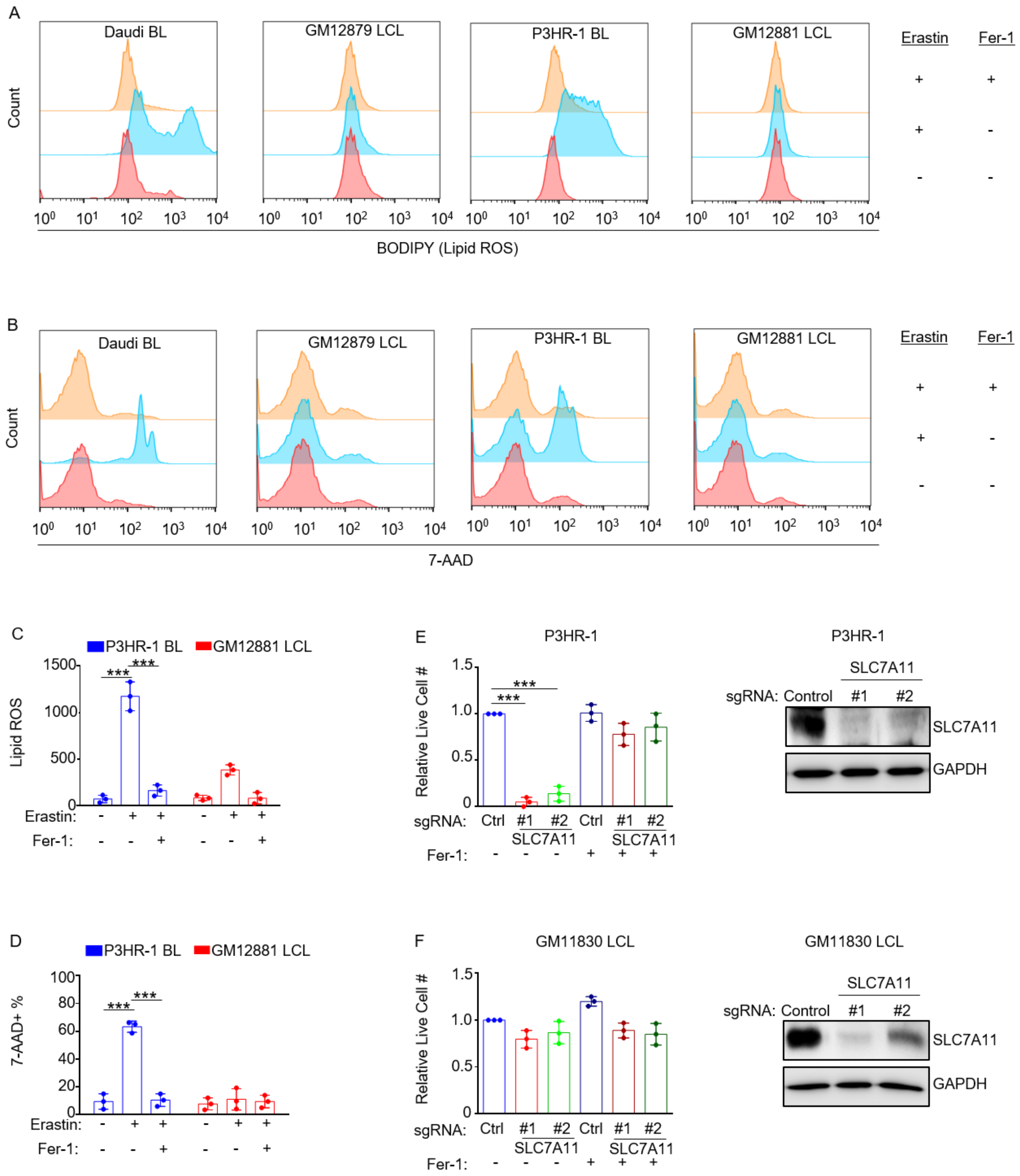


**Fig. S3. EBV+ Burkitt cells are more susceptible to SLC7A11 inhibition than LCLs.**

(A) Representative FACS histograms from n=3 replicates of BODIPY levels in Daudi or P3HR-1 BL cells versus GM12879 or GM12881 LCLs treated with erastin (10 µM) or Fer-1 (5 µM) for 18 hours.

(B) Representative FACS histograms from n=3 replicates of 7-AAD levels in Daudi or P3HR-1 BL cells versus GM12879 or GM12881 LCLs treated with erastin (10 μM) or Fer-1 (5 μM) for 24 hours.

(C) FACS mean + SEM lipid ROS levels from n=3 replicates of P3HR-1 or GM12881 cells treated with erastin or Fer-1 for 18 hours, as indicated.

(D) FACS %7-AAD mean + SEM levels from n=3 replicates of P3HR-1 or GM12881 cells treated with erastin or Fer-1 for 24 hours, as indicated.

(E-F) Relative mean + SEM live cell numbers from CellTitreGlo analysis from n=3 replicates of Cas9+ P3HR-1 BL (E) versus GM12878 LCL (F) after 7-days of control or SLC7A11 sgRNA expression, in the absence or presence of Fer-1 (5 μM), as indicated. Shown at right are representative immunoblots of WCL for SLC7A11 or GAPDH load control.

P-values were determined by one-sided Fisher's exact test. * $p<0.05$, ** $p<0.005$, *** $p<0.0005$.

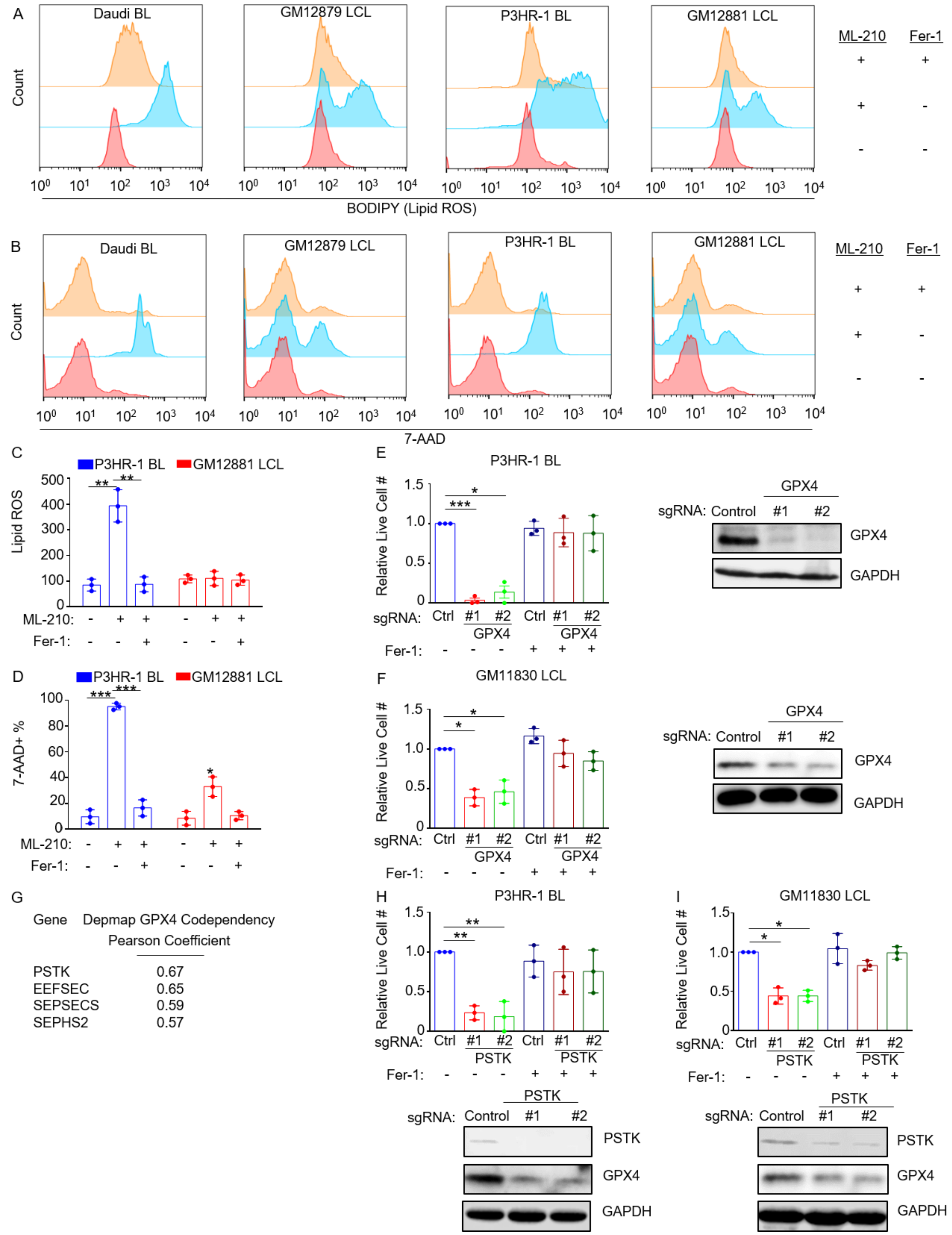


**Fig. S4: Susceptibility of EBV+ Burkitt versus LCLs to GPX4 inhibition.**

(A) Representative FACS histograms from n=3 replicates of BODIPY levels in Daudi or P3HR-1 BL cells versus GM12879 or GM12881 LCLs treated with ML-210 (1 μM) or Fer-1 (5 μM) for 18 hours.

(B) Representative FACS histograms from n=3 replicates of 7-AAD levels in Daudi or P3HR-1 BL cells versus GM12878 or GM12881 LCLs treated with ML-210 (1 μM) or Fer-1 (5 μM) for 24 hours.

(C) FACS mean + SEM lipid ROS levels from n=3 replicates of P3HR-1 or GM12881 cells treated with ML-210 or Fer-1 for 18 hours, as indicated.

(D) FACS %7-AAD mean + SEM levels from n=3 replicates of P3HR-1 or GM12881 cells treated with ML-210or Fer-1 for 24 hours, as indicated.

(E-F) Relative mean + SEM live cell numbers from CellTitreGlo analysis from n=3 replicates of Cas9+ P3HR-1 BL (E) versus GM11830 LCL (F) after 7-days of control or GPX4 sgRNA expression, in the absence or presence of Fer-1 (5 μM), as indicated. Shown at right are representative immunoblots of WCL for GPX4 or GAPDH load control.

(G) Broad Institute Dependency Map (DepMap) gene target with the highest Pearson coefficient of codependency with GPX4. High co-dependency indicates similar effects of gene knockout on target cell viability across the DepMap cell line collection of approximately 700 cell lines.

(H-I) Relative mean + SEM live cell numbers from CellTitreGlo analysis from n=3 replicates of Cas9+ P3HR-1 BL (E) versus GM11830 LCL (F) after 7-days of control or PSTK sgRNA expression, in the absence or presence of Fer-1 (5 μM), as indicated. Shown at right are representative immunoblots of WCL for PSTK or GAPDH load control.

P-values were determined by one-sided Fisher's exact test. * $p<0.05$, ** $p<0.005$, *** $p<0.0005$.

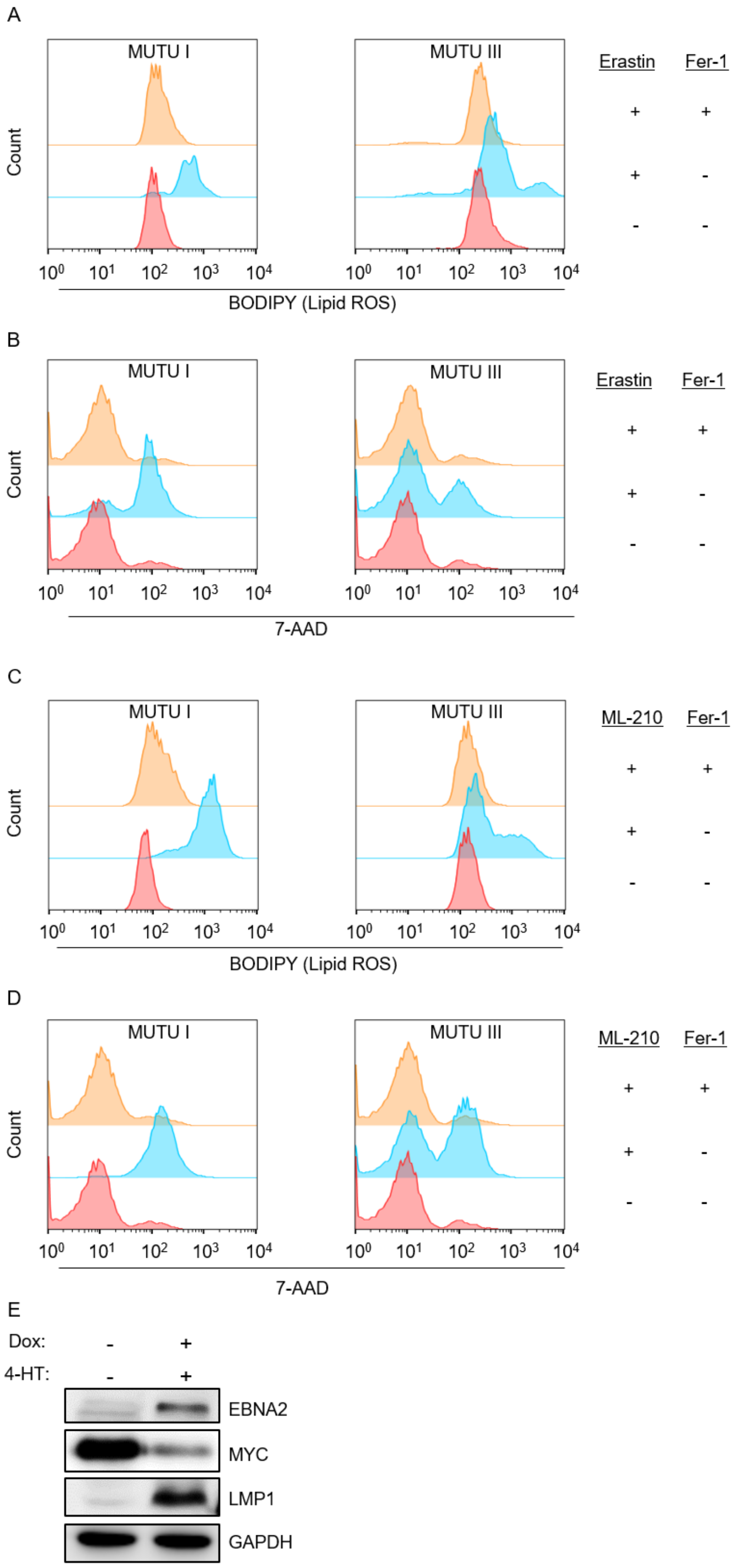
A
MUTU I
MUTU III
Count
Erastin
Fer-1
+ +
+ -
- -
$10^0$ $10^1$ $10^2$ $10^3$ $10^4$
BODIPY (Lipid ROS)
B
MUTU I
MUTU III
Count
Erastin
Fer-1
+ +
+ -
- -
$10^0$ $10^1$ $10^2$ $10^3$ $10^4$
7-AAD
C
MUTU I
MUTU III
Count
ML-210
Fer-1
+ +
+ -
- -
$10^0$ $10^1$ $10^2$ $10^3$ $10^4$
BODIPY (Lipid ROS)
D
MUTU I
MUTU III
Count
ML-210
Fer-1
+ +
+ -
- -
$10^0$ $10^1$ $10^2$ $10^3$ $10^4$
7-AAD
E
Dox: - +
4-HT: - +
EBNA2
MYC
LMP1
GAPDH

**Fig. S5. Differential effects of EBV latency I versus III on sensitivity to transformed B-cell ferroptosis induction.**

(A) Representative FACS histograms from n=3 replicates of BODIPY levels in MUTU 1 versus III BL cells treated with erastin (10 µM) or Fer-1 (5 µM) for 18 hours.

(B) Representative FACS histograms from n=3 replicates of 7-AAD levels in MUTU 1 versus III BL cells treated with erastin (10 µM) or Fer-1 (5 µM) for 24 hours.

(C) Representative FACS histograms from n=3 replicates of BODIPY levels in MUTU 1 versus III BL cells treated with ML-210 (1 µM) or Fer-1 (5 µM) for 18 hours.

(D) Representative FACS histograms from n=3 replicates of 7-AAD levels in MUTU 1 versus III BL cells treated with erastin (10 µM) or Fer-1 (5 µM) for 24 hours.

(E) Immunoblot of P-493 cell WCL from cells grown in the absence or presence of doxycycline (Dox, 1 µg/mL) and 4-hydroxytamoxifen (4HT, 2 µM) for three days.

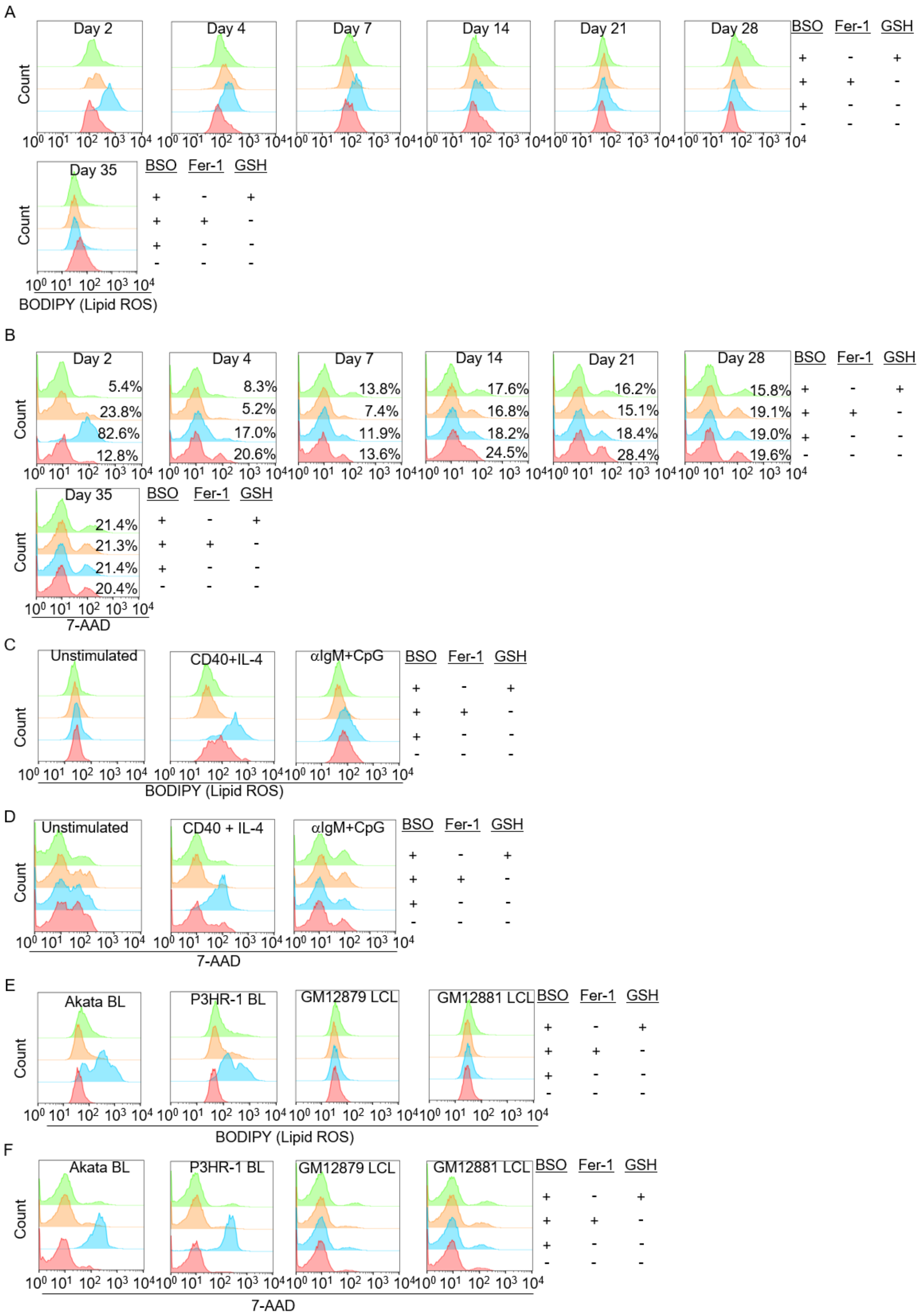
A
Day 2
Day 4
Day 7
Day 14
Day 21
Day 28
Day 35
BSO Fer-1 GSH
+ - +
+ + -
+ - -
- - -
Count
BODIPY (Lipid ROS)
B
Day 2
5.4%
23.8%
82.6%
12.8%
Day 4
8.3%
5.2%
17.0%
20.6%
Day 7
13.8%
7.4%
11.9%
13.6%
Day 14
17.6%
16.8%
18.2%
24.5%
Day 21
16.2%
15.1%
18.4%
28.4%
Day 28
15.8%
19.1%
19.0%
19.6%
Day 35
21.4%
21.3%
21.4%
20.4%
7-AAD
C
Unstimulated
CD40+IL-4
αIgM+CpG
BODIPY (Lipid ROS)
D
Unstimulated
CD40 + IL-4
αIgM+CpG
7-AAD
E
Akata BL
P3HR-1 BL
GM12879 LCL
GM12881 LCL
BODIPY (Lipid ROS)
F
Akata BL
P3HR-1 BL
GM12879 LCL
GM12881 LCL
7-AAD

**Fig. S6. Effects of GCLC inhibitor BSO on EBV-infected or stimulated B-cell lipid ROS and cell death.**

(A) Representative FACS histograms of BODIPY levels from n=3 replicates of primary B-cells infected with EBV for the indicated days and then treated with BSO (100 µM), Fer-1 (5 µM) or GSH (2.5 mM) for an additional 72 hours as indicated.

(B) Representative FACS histograms from n=3 replicates of 7-AAD levels in infected with EBV for the indicated days and then treated with BSO (100 µM), Fer-1 (5 µM) or GSH (2.5 mM) for an additional 72 hours as indicated.

(C) Representative FACS histograms of BODIPY levels from n=3 replicates of primary B-cells unstimulated or stimulated by Mega-CD40L (50 ng/mL) and IL-4 (20 ng/mL) or αIgM (1 µg/mL) and CpG (1 µM) as indicated for 2 days, and then treated with BSO (100 µM), Fer-1 (5 µM) or GSH (2.5 mM) under the same stimulation conditions for an additional 48 hours as indicated.

(D) Representative FACS histograms of 7-AAD levels from n=3 replicates of primary B-cells unstimulated or treated with Mega-CD40L (50 ng/mL) and IL-4 (20 ng/mL) or αIgM (1 µg/mL) and CpG (1 µM) as indicated for 2 days, and then treated with BSO (100 µM), Fer-1 (5 µM) or GSH (2.5 mM) under the same stimulation conditions for an additional 72 hours as indicated.

(E) Representative FACS histograms from n=3 replicates of BODIPY levels in Akata or P3HR-1 BL versus GM12879 or GM12881 LCLs cells treated with BSO (100 µM), Fer-1 (5 µM) or GSH (2.5 mM) for 48 hours.

(F) Representative FACS histograms from n=3 replicates of 7-AAD levels in Akata or P3HR-1 BL versus GM12879 or GM12881 LCLs cells treated with BSO (100 µM), Fer-1 (5 µM) or GSH (2.5 mM) for 48 hours.

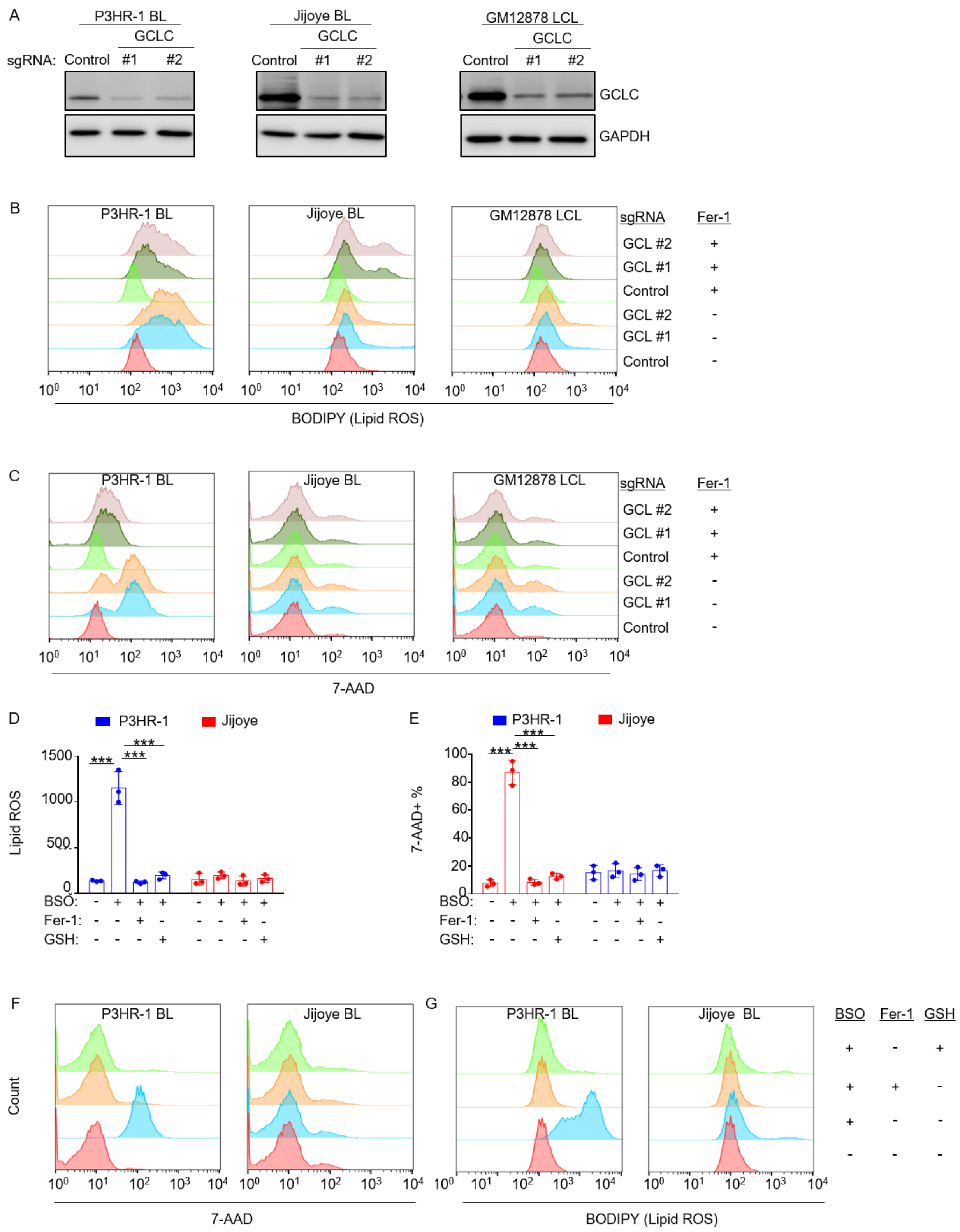


**Fig. S7. Chemical or genetic perturbation kills latency I, but not latency III, EBV infected cell lines**

(A) Representative immunoblot of WCL from n=3 replicates of Cas9+ P3HR-1, Jijoye or GM12878 expressing control or GCLC sgRNAs for 5 days.

(B) Representative FACS histograms from n=3 replicates of BODIPY levels in Cas9+ P3HR-1, Jijoye or GM12878 expressing control or GCLC sgRNAs for 12 days and treated with Fer-1 (5 μM) as indicated.

(C) Representative FACS histograms from n=3 replicates of 7-AAD levels in Cas9+ P3HR-1, Jijoye or GM12878 expressing control or GCLC sgRNAs for 14 days and treated with Fer-1 (5 μM) as indicated.

(D) FACS BODIPY mean + SEM values from n=3 replicates of P3HR-1 or Jijoye BLs treated with BSO (100 μM), Fer-1 (5 μM) or GSH (2.5 mM) as indicated for 48 hours.

(E) FACS &7-AAD+ mean + SEM values from n=3 replicates of P3HR-1 or Jijoye BLs treated with BSO (100 μM), Fer-1 (5 μM) or GSH (2.5 mM) as indicated for 72 hours.

(F) Representative FACS histograms of 7-AAD levels from n=3 replicates of P3HR-1 or Jijoye BL treated with BSO (100 μM), Fer-1 (5 μM) or GSH (2.5 mM) as indicated for 72 hours.

(G) Representative FACS histograms of Bodipy levels from n=3 replicates of P3HR-1 or Jijoye BL treated with BSO (100 μM), Fer-1 (5 μM) or GSH (2.5 mM) as indicated for 48 hours.

P-values were determined by one-sided Fisher's exact test. * $p<0.05$, ** $p<0.005$, *** $p<0.0005$.

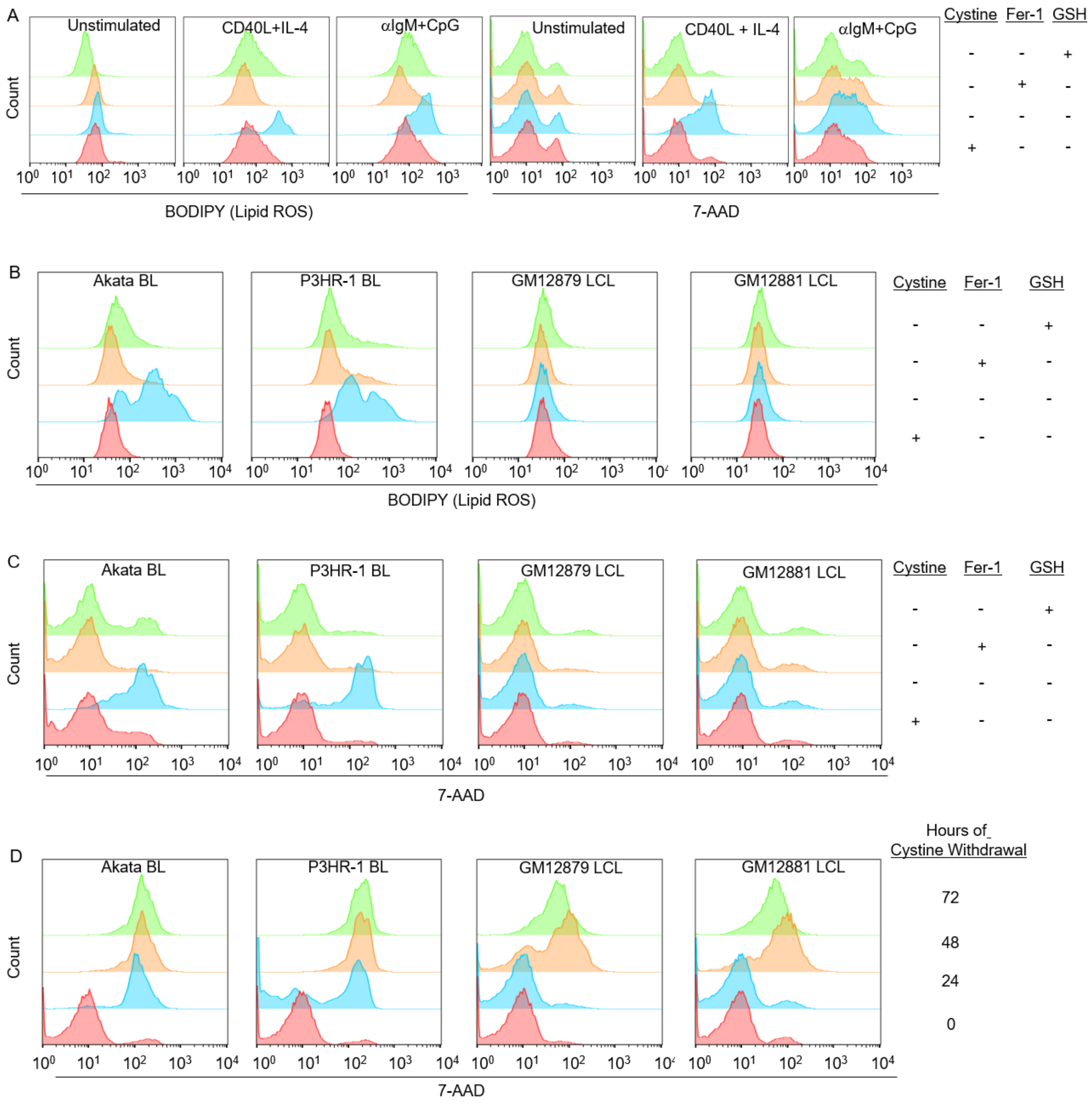


**Fig. S8. Effects of cystine withdrawal on EBV-infected vs stimulated B-cell ferroptosis.**

(A) Representative FACS histograms from n=3 replicates of BODIPY (left) or 7-AAD (right) levels in primary B-cells unstimulated or stimulated by Mega-CD40L (50 ng/mL) and IL-4 (20 ng/mL) or αIgM (1 µg/mL) and CpG (1 µM) as indicated for 2 days, and then again for another 2 days for BODIPY analysis or 3 days for 7-AAD analysis in media with or without cystine, Fer-1 or GSH, as indicated.

(B) Representative FACS histograms from n=3 replicates of BODIPY levels in Akata or P3HR-1 BL or GM12879 or GM12881 LCL grown in media without or with cystine, Fer-1 (5 µM) or GSH (2.5 mM) for 48 hours.

(C) Representative FACS histograms from n=3 replicates of 7-AAD levels in Akata or P3HR-1 BL or GM12879 or GM12881 LCL grown in media without or with cystine, Fer-1 (5 μM) or GSH (2.5 mM) for 72 hours.

(D) Representative FACS plots of 7-AAD levels from n=3 replicates of EBV-Akata and P3HR-1 BL or GM12879 and GM12881 LCL seeded into media without cystine for the indicted times.

P-values were determined by one-sided Fisher's exact test. * p, <0.05, **p<0.005, ***p<0.0005.

#### Supplementary Methods

**Cell culture.** 293T, Daudi and Jijoye were purchased from American Type Culture Collection and cultured in DMEM with 10% fetal bovine serum (FBS, Gibco) and 1% penicillin-streptomycin (Gibco) in a humidified incubator at 37°C and 5% CO2. GM11830, GM12878, GM12879, and GM12881 LCL were obtained from Coriell. MUTU I, MUTU III, Kem I, Kem III were kind gifts from Jeff Sample and Alan Rickinson. All B-cell lines were cultured in RPMI-1640 (Invitrogen) supplemented with 10% standard FBS and penicillin-streptomycin in a humidified incubator at 37°C and at 5% CO2. P493-6 LCLs were obtained from Micah Luftig. P493-6 cells have a conditional EBNA2-HT allele and also a Tet-off exogenous *MYC* allele. In the absence of tetracyclines, exogenous MYC is highly induced. In the presence of 4HT and presence of tetracyclines, P-493 grow as LCLs. P493-6 cells were maintained in the absence of 4HT or doxycycline to induce a BL-like state of high MYC expression. To grow in the lymphoblastoid cell state (intermediate MYC, EBV latency III), P493-6 cells were grown in the presence of both 1 μM 4HT and 1 μM doxycycline. After 72 hours of growth in any of these conditions, cells were collected and treated as described. All cells were routinely confirmed to be mycoplasma-negative by Lonza MycoAlert assay (Lonza).

**Antibodies and reagents.** Antibodies against the following proteins were used in this study: GPX4 (Proteintech, Cat#14432), PSTK (Genetex, Cat#GTX45610), GAPDH (EMD Millipore, Cat#MAB374), rabbit anti-human MYC (Santa Cruz Biotechnology, sc-764), PE anti-CD23 (BD, Cat#555711), rat anti-EBNA2 (EMD Millipore, Cat#MABE8), mouse anti-LMP1 (Abcam, Cat#ab78113). The following chemicals were obtained from Sigma-Aldrich: ferrostatin-1 (Cat#SML0583-25MG), erastin (Cat#E7781-5MG), buthionine sulfoximine (BSO) (Cat#B2515-500MG), L-glutathione, reduced (Cat#G6013-5G). ML-210 (Cat#23282) was obtained from

Cayman Chemicals. H21491). Mega-CD40L was purchased from Enzo (ALX-522-110-C010) and CpG oligonucleotide from Invivogen. Anti-IgM was purchased from Sigma-Aldrich (I0759-5X1MG).

**Primary human B-cell isolation and culture.** Platelet-depleted venous blood obtained from the Brigham & Women's hospital blood bank were used for primary human B cell isolation, following our Institutional Review Board-approved protocol for discarded and de-identified samples. Our studies on primary human blood cells were approved by the Brigham & Women's Hospital Institutional Review Board. For most experiments, cells were cultured in RPMI-1640 (Invitrogen) supplemented with 10% standard FBS and penicillin-streptomycin. Cells were cultured in a humidified incubator at 37$^{o}$C and at 5% CO2.

**Cystine Depletion.** Primary B cells and B cell lines were collected and washed three times with 1x PBS supplemented with 0.5% Bovine serum albumin (BSA, Sigma Aldrich Cat#A7906-100G). Afterwards, cells were reseeded at 5x10^5 cells / mL in RPMI 1640 (MP Biomedicals, Cat# ICN1646454) supplemented with dialyzed FBS (Gibco, Cat# 26400044) replete with L-cystine (Sigma Aldrich, Cat#,C2526-100G), L-methionine (Sigma Aldrich, Cat# M5308-25G) and L-glutamine (Gibco, Cat# 25030-164) as a control, or without L-cystine. Cells were harvested at indicated timepoints for lipid peroxide quantification using BODIPY C-11 or viability using 7-AAD.

**Immunoblot analysis.** Cells were lysed in Laemelli buffer (0.0625 M Tris Base, 0.07M SDS, 10% glycerol v/v, 5% 2-mercaptoethanol, .002% bromophenol blue) and sonicated at 4 degrees C for 10 seconds using a probe sonicator at the maximum setting. Lysates were centrifuged at 17,000 x g for 30 minutes and supernatant boiled for 10 minutes. SDS-PAGE was performed using 10 or 14% acrylamide gels and transferred onto nitrocellulose membranes for 80 minutes at 100 V at 4˚C. Membranes were blocked for 1 hour at room temperature in 5% milk / 1x TBS-T, probed with primary antibodies (diluted in 1x TBS-T with 0.02% sodium azide at recommended manufacturer concentrations) overnight at 4˚C, followed by a one hour incubation with respective HRP-conjugated secondary antibodies in 1x TBS-T. Blots were imaged on Li-Cor Odyssey workstation.

**Flow cytometry.** Cells were harvested and washed twice with PBS/2% FBS. In order to account for differences in background viability produced by uninfected cells, we stained cells collected during the first four days post infection using a CD23 antibody to demarcate EBV infected cells from uninfected, a known marker of EBV infection (1, 2). Cells were then incubated with a 1:200 dilution of PE-conjugated mouse anti-CD23 antibody (BD, Cat#555711) antibody in PBS supplemented with 2% FBS for 30 minutes at room temperature, protected from light. Cells were washed twice with PBS supplemented with 2% FBS and then resuspended in 200 μL of PBS supplemented with 2% FBS and analyzed immediately via flow cytometry. For 7-AAD (Thermo Fisher, Cat#A1310) viability assays, cells were harvested and washed twice with 1x PBS supplemented with 2% FBS (Gibco). Cells were then incubated with a 1 μg / mL 7-AAD solution in 1x PBS / 2% FBS for five minutes at room temperature, protected from light. Cells were then analyzed via flow cytometry. For lipid peroxidation assays, 10^5 cells were harvested, centrifuged (300 g x 5 minutes) and incubated in a u-bottom 96-well plate in 200 μL BODIPY C-11 581/591 working solution (2.5 μM BODIPY in RPMI 1640 supplemented with 10% FBS). Cells were then incubated at 37°C with 5% $CO_2$ for 30 minutes, protected from light. Cells were then washed twice with 1x PBS supplemented with 2% FBS before immediate analysis via flow cytometry For CFSE labeling and cell proliferation assays, CellTrace™ CFSE solution was prepared according to manufacturer's instructions. 10^7 primary B cells were resuspended and incubated in one mL of CellTrace™ working solution for ten minutes in a 37°C / 5% CO2 incubator, protected from light. Taking care to protect cells from light as much as possible, five mL of RPMI 1640 with 10% FBS was added to the stained cells. Cells were incubated at room temperature for five minutes, protected from light, to remove free dye and prevent toxicity. Cells were then pelleted by centrifugation (300g x 5 minutes, room temperature) and resuspended in fresh RPMI with 10% FBS three times before the cells were resuspended at a ferroptosis inducing agental concentration of one million cells / mL of fresh RPMI with 10% FBS. Cells were then stimulated or infected as indicated. CellTrace™ Violet solution was prepared according to manufacturer's instructions. To make a working solution, all of the resuspended CellTrace™ Violet was diluted in 10 mL of 1x PBS warmed to 37 degrees Celsius. 5x10^7 primary B cells were resuspended in one mL of this CellTrace™ Violet working solution and incubated in a 37 degree Celsius incubator for 20 minutes, protected from light. After 20 minutes, 5 mL of RPMI (supplemented with 10% FBS) was added directly to the labeled cells and cells were incubated for another 5 minutes at 37 degree Celsius to quench the CellTrace™ Violet dye. Cells were then centrifuged as above and stimulated as indicated. In order to analyze cell proliferation, labeled cells were analyzed via flow cytometry with a BD FACSCalibur instrument and analysis was performed with FlowJo V10.

**CRISPR/Cas9 mutagenesis.** B-cell lines with stable Cas9 expression were established as described previously (3). sgRNA constructs were generated as previously described (4) using sgRNA sequences from the Broad Institute Avana or Brunello libraries. CRISPR editing was performed as previously described (4). Briefly, lentiviruses encoding sgRNAs were generated by transient transfection of 293T cells with packaging plasmids and pLentiGuide-Puro plasmids. Daudi, MUTU I, GM12878, and GM11830 cells stably expressing Cas9 were transduced with the lentiviruses and selected with 3 μg/mL puromycin for three days before replacement with antibiotic-free media. CRISPR editing was confirmed by immunoblotting 3 days post puromycin selection. The sgRNAs used in this study were constructed using oligos based on the sequences below:

| sgRNA name | Sequence |
|---|---|
| GPX4 #1 | TAACCTGGACAAGTACCGGT |
| GPX4 #2 | GTGCCCGTCGATGTCCTTGG |
| PSTK #1 | CGTCCTCTGTGGCCTCCCCG |
| PSTK #2 | GGTTCCTCTGATGTTCTCGG |
| GCLC #1 | AGGCCAACATGCGAAAACGC |
| GCLC #2 | AGGCCAACATGCGAAAACGC |
| SLC7A11 #1 | AAGGGCGTGCTCCAGAACAC |
| SLC7A11 #2 | GAAGAGATTCAAGTATTACG |

**Table S1: Table of sgRNAs used to knockout genes of interest.**

sgRNAs used for CRISPR-Cas9-mediated knockout of the indicated genes are listed above.
